# Nandrolone alters the behavioral response to cocaine as well as striatal and cortical dopamine receptors of prepubertal male rats

**DOI:** 10.1101/2024.02.07.579337

**Authors:** Jaime A. Freire-Arvelo, Carlos J. Rivero, Iván G. Santiago-Marrero, Andrea Mendez-Morales, Amanda J González-Segarra, Enrique U. Pérez-Cardona, Ricardo J. Torres-Ramirez, Annabell C. Segarra

## Abstract

Nandrolone is the anabolic androgenic steroid (AAS) most used by athletes and adolescents. The use of supraphysiologic doses has been associated with dysfunction in brain areas that regulate anxiety, motivation, and reward. This study investigated if exposure to nandrolone before puberty altered anxiety- and addictive-like behaviors. Changes in dopamine type 2 receptors (D2DR) in the nucleus accumbens (NAc) and medial prefrontal cortex (mPFC) were also examined. Male rats received 10 daily injections of nandrolone decanoate (20 mg/kg) starting on day 28. Afterwards, they were tested in the elevated plus maze (EPM) and open field (OF). Their locomotor response (sensitization) and preference (conditioned place preference (CPP)) to cocaine (15 mg/kg) was also assessed. Nandrolone reduced anxiety and ambulation. Nandrolone-treated males also displayed sensitization to cocaine at an earlier age (day 44) than oil-treated males (day 52) and showed a 27% reduction in CPP to cocaine. Expression of D2DR in the NAc, and in the PFC of males tested for CPP was increased by nandrolone, whereas treatment with cocaine reduced accumbal D2DR. We hypothesize that nandrolone accelerates the development of the neural circuitry that regulates behavioral sensitization and reduces the rewarding property of cocaine, as manifested in the reduction of CPP. It is possible that the observed increase in accumbal D2DR may partially mediate the reduced anxiety and ambulation as well accelerate the maturation of the neural circuitry responsible for the sensitized response to cocaine.

## Introduction

Anabolic-androgenic steroids (AAS) are derivatives of the main male gonadal hormone testosterone (T) developed in the 1950s to treat men with hypogonadism, delayed puberty (Basaria and Dobs, 2001) and to promote growth. The term anabolic comes from their effects of promoting metabolic processes, such as protein synthesis in muscle tissue and erythropoiesis (Kochakian and Welder, 1993). Androgenic comes from their masculinizing properties. Their anabolic properties make these substances appealing to athletes. AAS provides athletes with an edge in training and in competition, they increase muscle strength and endurance and promote competitive and aggressive behaviors (Bahrke et al., 1996).

Before the ’90s, the use of AAS was circumscribed mainly to athletes during training and before competitions (Sjöqvist et al., 2008). However, during the last two decades, young men have been using AAS all year round to improve their physical appearance (Denham, 2006; Kindlundh et al., 1999). Use of AAS as self-medication is also on the rise among the female-to-male transgender population (Metastasio et al., 2018). In the USA, about 3-4 million people have used AAS, of those approximately 1 million have developed dependency (Pope et al., 2014a). Worldwide prevalence of AAS use is 3.3%, being higher for males (6.4%) than for females (1.6%) (Sagoe et al., 2014). Users of AAS report administering dosages that can be higher than 100X the physiological dose (Brower et al., 1990). This is of significant concern because AAS have deleterious side effects, particularly on the cardiovascular system. AAS also cause hepatic toxicity, decrease fertility and alter secondary sexual characteristics (Parkinson and Evans, 2006; van Amsterdam et al., 2010). Many studies find that AAS increase psychiatric disorders (Pope et al 2014b; Oberlander and Henderson, 2012; Onakomaiya and Henderson, 2016). Some of the symptoms are associated with the exposure to AAS (mania, aggression, risk-taking behaviors, irritability) whereas others can result from AAS withdrawal (depression, loss of libido, suicidal thoughts, hypersomnia) (Pagonis et al., 2006; Pope and Katz, 1988). There is considerable variability in the display and range of these symptoms, in very few individuals these symptoms are disabling (Kaufman et al., 2015).

Androgens, including AAS, have rewarding properties (Johnson and Wood, 2001; Wood, 2004, 2002) and may contribute to the development of substance abuse and dependency to other drugs of abuse (Brower et al., 1989). Approximately 30-32% of subjects using AAS will develop a dependency to AAS (Pope et al., 2014a), this risk is higher for women and adolescents of both sexes (Kanayama et al., 2008; Penatti et al., 2011). AAS also promote dependence on other drugs such as opioids and cocaine (Pope et al., 2014a). Several studies report that abuse of drugs, particularly of cocaine, by AAS users is higher than in the general population (DuRant et al., 1995; A. M S Kindlundh et al., 2001). Because of their widespread use and adverse side effects, AAS were classified as a schedule III drug by the Drug Enforcement Agency (DEA).

Among the AAS, nandrolone (19-nortestosterone), in its long-lasting ester form of nandrolone decanoate (ND), is the AAS most widely used worldwide (Lood et al., 2012). The androgenic activity of this compound is lower than that of dihydrotestosterone (DHT); in contrast, its anabolic properties are higher than those of T (Saartok et al 1984) making it attractive for abuse by male and female athletes. Nandrolone has a higher affinity for the androgen receptor (AR) than T and is less susceptible to degradation by the enzyme 17 beta-hydroxysteroid dehydrogenase, which may explain its enhanced anabolic properties. Moreover, although nandrolone and T can be reduced to DHT in target tissue that contains the enzyme 5 alpha-reductase, enzyme binding to nandrolone is weaker than to T (Bergink et al., 1985; Kicman, 2008; van der Vies et al., 1993). Also, binding of nandrolone to the AR is weaker than that of DHT. Taken together, this explains the stronger effects of nandrolone compared to T, on target tissues with no 5 alpha-reductase activity, and the weaker effect on tissues with a high 5 alpha-reductase activity, resulting in greater anabolic/androgenic properties (Tenniswood et al., 1982; van der Vies et al., 1985).

Adolescents, compared to children and adults, show the highest incidence of risk-taking behavior and of experimentation with drugs of abuse (SAMHSA, 2019). AAS are among the drugs that are most used by adolescents, particularly males. This is a significant concern since AAS can cross-sensitize with other drugs of abuse. Cocaine, one of the primary drugs used by people that abuse AAS, shares many of the harmful side effects on the cardiovascular system and also promotes risk-taking behavior, aggravating the detrimental health consequences when using both drugs.

Drugs of abuse exert their behavioral and addictive effects by acting on certain areas of the brain associated with decision making, motivation and reward, such as the prefrontal cortex (PFC) and nucleus accumbens (NAc) (Baker et al. 2003).

Dopaminergic receptors in these brain areas participate in regulating addictive behaviors (Beyer and Steketee, 2002; Steketee and Walsh, 2005). In particular, decreased D2-like dopamine receptors (D2DR) expression in these brain areas are associated with increased novelty seeking and risk-taking, behaviors associated with addiction (Goldstein and Volkow, 2002; Koob and Volkow, 2010). Manipulating striatal D2DR can alter the response to drugs of abuse, such as cocaine. Mice that lack striatal presynaptic D2DR show increased sensitivity to the locomotor activating effects of cocaine (Bello et al, 2011), whereas rats with increased D2DR sensitivity show enhanced cocaine self-administration (Marinelli and White, 2000).

This study investigated if exposure to nandrolone prior to puberty affected anxiety-like behaviors as well as the behavioral response to cocaine. The nucleus accumbens and prefrontal cortex were also examined to determine if nandrolone induces changes in the D2DR population of these brain substrates. To accomplish these goals, rats were injected with nandrolone decanoate (20 mg/kg) during postnatal days (PN) 28-37 and their behaviors assessed from days 38 to 62. Animals were sacrificed at days 54 or 63 and brain tissue removed for further analysis.

## 2. Materials and Methods

**2.1 Animals**

Pregnant Sprague Dawley rats were purchased from Charles Rivers Laboratories (Ballardvale St, MA, USA). Dams were housed in pairs, with water and Purina® rat chow provided ad libitum. They were maintained in a temperature and humidity-controlled room, under a (12L:12D) light-dark cycle with lights off at 5 PM. After parturition, pups were cross fostered, and each dam was housed separately with their litter. The litter was culled to 10 pups per dam, 5 males and 5 females. Pups remained with the mother until weaning (day 23). All animal experiments were approved by the Institutional Animal Care and Use Committee (IACUC) of the University of Puerto Rico, Medical Sciences Campus (Protocol 1140215) and adhere to USDA, NIH and AAALAC guidelines.

### 2.2 Drugs and chemicals

The AAS used in this study was 4-estren-17beta-ol-3-one decanoate (nandrolone decanoate) (Steraloids, Inc., Newport, RI, USA), dissolved in sesame oil. Nandrolone was administered subcutaneously (s.c.) at a dose of 20 mg/kg. The nandrolone dose of 20 mg/kg was selected since it is similar to the supraphysiological doses used by AAS users (Clark and Henderson, 2003). Cocaine-HCl (Sigma-Aldrich, St. Louis, MO, USA) was dissolved in 0.9% sterile saline and administered intraperitoneally (i.p.) at a dose of 15 mg/kg. The dose of 15 mg/kg of cocaine has been used extensively in our laboratory and has been proven effective in inducing behavioral sensitization and CPP to cocaine. We have found that higher doses (30 mg/kg) may induce tolerance and lower doses are not as effective in a behavioral sensitization paradigm that is not context-dependent (Segarra et al., 2014; Menendez-Delmestre and Segarra, 2011).

### 2.3 Nandrolone Treatment

At postnatal day 28 (PN-28) rats were weighed and randomly distributed into two groups: one that received daily injections of nandrolone (ND) (20mg/kg) (n=20) and the other that received daily injections of sesame oil (Oil) (n=20) for 10 consecutive days. Two cohorts of rats were used for the behavioral studies. One cohort was used for the Open Field Test (OFT) and the sensitization experiments. The second cohort was used for the Elevated Plus Maze (EPM) studies and the CPP experiments. On day 39 one cohort of rats were tested in the Open Field (OFT). From day 40 to 62 they were tested for behavioral sensitization to cocaine and euthanized on day 63. The second cohort of rats were tested on day 39 on the Elevated Plus Maze (EPM) and from days 40 to 53 they were used to assess Conditioned Place Preference (CPP) to cocaine. This last group of rats were euthanized on day 54. For the CPP and Sensitization tests, animals were assigned to one of 4 groups: Oil-Saline (Oil-Sal), Oil-Cocaine (Oil-Coc), Nandrolone-Saline (ND-Sal), Nandrolone-Cocaine (ND-Coc).

### 2.4 Elevated Plus Maze

The EPM is a highly used paradigm to measure anxiety-related behaviors (Pellow et al., 1985). Rats previously exposed to anxiogenic drugs decrease the time spent in the open arms of the maze, while anxiolytic drugs increase the time spent in open arms (Biedermann et al., 2017). Our testing apparatus consisted of a plus-shaped custom-made apparatus with two 50 cm open arms and two 50 cm enclosed arms, each with an open roof. The floor of the open arms had a 1 cm ledge to prevent rats from slipping to the floor. We also lined the floor with rugged plastic to avoid slipping. The apparatus was elevated 70 cm from the floor. An infrared video camera was placed in the center above the maze. The camera was connected to a computer containing the ANY-Maze™ software. At the beginning of the test, rats were placed at the junction of the open and closed arms and the video tracking system was activated. The software automatically records the number of entries into the open and closed arms, as well as of the time spent in each arm. Entry into an arm was defined as the time point when more than 95% of the rat is in the arm. This was considered time zero. The test ended after 5 min. The amount of time spent in the closed arms, the time spent in the open arms relative to the closed arms and the number of entries into the open arms was measured. The greater the amount of time spent in the closed arms, the higher the anxiety.

### 2.5 Open field test

The OFT is an assay of locomotor activity that can be used to measure anxiety, exploratory behavior, risk taking behavior and thigmotaxis. Anxiolytic drugs increase the amount of time rats spend in the center area (Treit and Fundytus, 1988). The locomotor activity chambers (Versamax™) were used to measure Open Field Behavior. These chambers are made from clear acrylic (42 cm × 42 cm × 30 cm). All beams were connected to a Data Analyzer that sent information to a personal computer.

Animals were placed in the Activity Cage and allowed to roam freely for 10 minutes. The breaking of infrared beams determined the position of the rats in the activity cage. The amount of time spent in the center of the cage versus at the periphery was compared, as well as the total distance travelled. Animals that spent less time in the center of the cage were classified as more anxious and as displaying less risk-taking behavior when compared to their counterpart controls (oil-treated).

### 2.6 Locomotor activity

Horizontal and stereotyped activity were measured with the automated animal activity cage system (Versamax™; AccuScan Instruments, Columbus, Ohio). The cages were made from clear acrylic (42 cm × 42 cm × 30 cm) with 16 equally spaced (2.5 cm) infrared beams across the length and width of the cage at a height of 2 cm (horizontal beams). An additional set of 16 infrared beams were located at a height of 10 cm (stereotyped activity). This system differentiates between horizontal, stereotyped, or rearing activity based on sequential breaking of different horizontal beams, the same beams or vertical beams respectively.

Activity was measured in an isolated room with low illumination. Animals were habituated to the chamber for 1 hour, 1 day prior to injections (Day 0). On days 1,5,13 and 23 rats were placed for 30 mins in the chambers, and basal locomotor activity was recorded. Animals then received a saline or cocaine injection, and locomotor activity was recorded for 60 additional minutes. On days 2–4 animals received a daily injection of 0.9% saline or cocaine (15mg/kg) in their home cages. During days 6–12 and 14-22 animals remained in their home cages undisturbed (Fig 1). Animals were sacrificed the day after the last behavioral test at 63 days of age.

**Figure 1.**
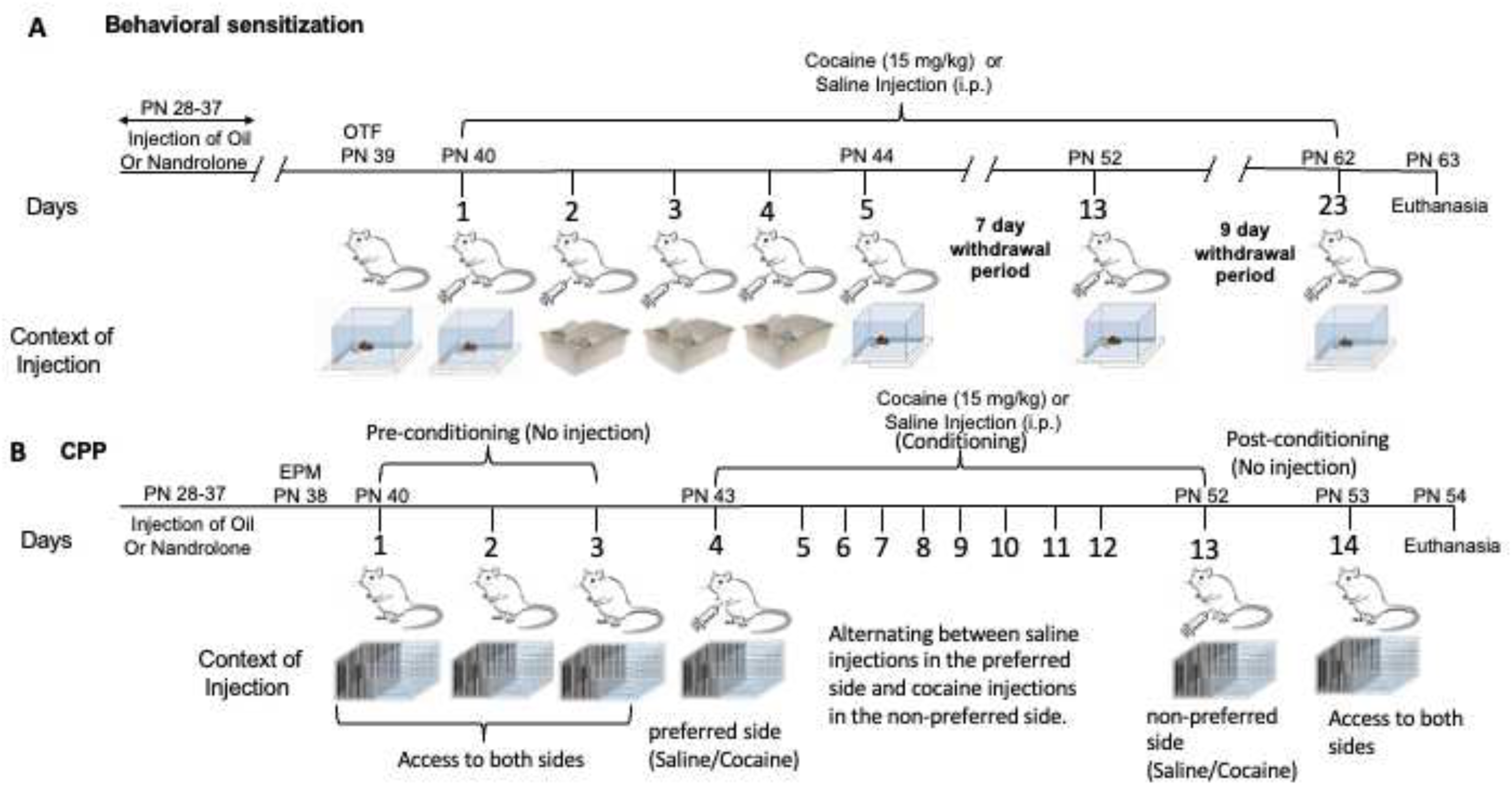
Behavioral sensitization (A) and CPP (B) protocols. **Fig 1A**. Five days after rats were weaned from their mother, they received a daily oil or nandrolone injection for ten consecutive days (PN-28 to 37). For the behavioral sensitization experiments, rats were habituated to the locomotor activity chamber for 1 hour (2 days post-nandrolone day, PN-39). The data obtained from the first ten minutes of habituation (Day 0) and of Day 1 was used as data for the open field test (Figs 2B and Fig 2C). From days 1 through 5, on day 13 and on day 23, rats were injected with saline or with cocaine (15 mg/kg). Rats were injected in the locomotor activity chamber (days 1, 5, 13 and 23) or in their home cage (days 2, 3 and 4) (see the context of injection). From days 6 through 12 and from days 14 through day 22, rats remained undisturbed in their home cages. **Fig 1B**. Five days after rats were weaned from their mother, they received a daily oil or nandrolone injection for ten consecutive days (PN-28 to 37). On day 2 (PN-38) a second group of rats were tested in the elevated plus maze (EPM). These same rats were used for the CPP experiments. To determine the rat’s preference for a particular CPP chamber, rats were allowed to roam freely through both chambers for 3 days (pre-conditioning). The amount of time spent in each chamber was calculated to determine which side it preferred. The following 10 days (days 4–13, conditioning days) rats were injected daily alternating between saline and cocaine (15 mg/kg) injections. Rats received cocaine in the non-preferred chamber and saline in the preferred chamber. Saline animals received saline in both chambers. After the injection, rats were confined for 30 min to the chamber where they received the injection. The last day (day 14, post-conditioning day), the animals were placed in the activity chamber and allowed to roam freely between the two chambers for 30 minutes. The amount of time spent in each chamber was recorded and compared to that of pre-conditioning.

### 2.6 Conditioned Place Preference

Cocaine-induced CPP was measured using Versamax™ activity cages. Each cage consisted of an acrylic box divided into 2 chambers and placed in the locomotor activity apparatus. For the pre- and post-conditioning sessions, the chambers were separated by an acrylic wall that had an opening; during the conditioning phase, a solid acrylic wall separated the two chambers. Each chamber had different visual and tactile cues. During pre-conditioning, animals were placed in the CPP apparatus for 3 consecutive days and allowed to roam between both chambers for 15 minutes. The amount of time spent in each chamber was recorded. The conditioning phase consisted of alternating saline and cocaine injections for 10 days. Saline was injected in the preferred chamber, cocaine in the non-preferred chamber with 24 hours of separation between injections. Rats were confined to the chamber where they received the injection for 30 min. During post-conditioning, animals were placed in the activity chamber and allowed to roam between the 2 chambers for 15 min (Fig 1). The time spent in each chamber during pre- and post-conditioning was compared. Rats that showed a significant increase in the time spent in the chamber where they received cocaine displayed conditioned place preference.

### 2.7 Western Blots

Western blots were used to quantify D2DR levels in the mPFC and NAc. The protein concentration of the samples was determined by the BioRad Protein Assay method (BioRad Laboratories, Hercules, CA, USA). For our gels, 20 μg of protein were mixed in SDS/-mercaptoethanol, vortexed, and heated at 95°C for 7 min prior to separation by 10% SDS-PAGE (BioRad Laboratories). Following electrophoresis, proteins were transferred to a 0.2 μm nitrocellulose membrane using a Trans-Blot Turbo (BioRad Laboratories). Nonspecific binding to the membrane was blocked by incubating in Odyssey Blocking Buffer for 60 min at room temperature. This was followed by overnight incubation at 4°C with a D2 receptor antibody (1:200; Santa Cruz Biotech, Santa Cruz, CA, USA, #sc-5303) and a B-Actin antibody (1:2500; Abcam, MA, #ab8227) dissolved in Odyssey Blocking Buffer. The next day, membranes were washed 3X in TRIS-buffered saline and Polysorbate 20 (PBS-T). After the washes, membranes were incubated for one hour in IRDye 680RD goat anti-rabbit (1:15000; LI-COR, Lincoln, NE, USA, #926-68071) and IRDye 800CW goat anti-mouse (1:15000; LICOR, Lincoln, NE, USA, #926-32210). Proteins were detected using the Odyssey CLx infrared imaging system (excitation/emission filters at 700 nm/ 800 nm range, LI-COR Biosciences, Lincoln, NE, USA). Optical density of D2 receptors of each sample was obtained using Odyssey software (LI-COR Biosciences), normalized against background, and then corrected against their own B-Actin levels.

## 3. Statistical analyses

All data were analyzed using GraphPad Prism version 9.00 for Windows (Graphpad Software, San Diego California USA). An unpaired t-test was used to compare two groups, a Two-way ANOVA was used when comparing more than two groups and a repeated measures MANOVA was used for analyzing repeated measures. To ascertain if rats showed sensitization, the time course of each group was analyzed separately using repeated measures (RM) ANOVA with days (40, 44, 52 and 62) and minutes (35-90) as the repeated factors. Tukey’s multiple comparisons were used for post-hoc analysis to compare locomotor and stereotyped activity across time. CPP was analyzed using repeated measures MANOVA, using Pre and Post Conditioning as repeated factors. A Two-Way ANOVA was used to compare D2DR expression in the PFC and the NAc between groups. Results of statistical analysis are included as supplemental material. Data are presented as the mean ± standard error of the mean (SEM). A p-value of less than 0.05 (p<0.05) was considered statistically significant.

## 4. Results

### 4.1 Open Field and basal locomotor activity

Rats treated with nandrolone spent more time in the center of the open field, and ambulated less, than oil-treated rats (Fig 2A and 2B respectively). Data were analyzed with Student’s T-test - Fig 2A: t=2.361, df=36, p = 0.0264; Fig 2B: t=5.174, df=36, p<0.0001. The decrease in distance travelled corresponds to the first 10 minutes of habituation (sensitization protocol-Day 0). A decrease in total horizontal activity during the first 10 min was also observed in the habituation portion on day 1 of the sensitization protocol (Fig 1C; Student’s T-test: t=2.152, df=18, p<0.0453). This difference was not observed on subsequent testing days (days 5, 13 and 23).

**Figure 2.**
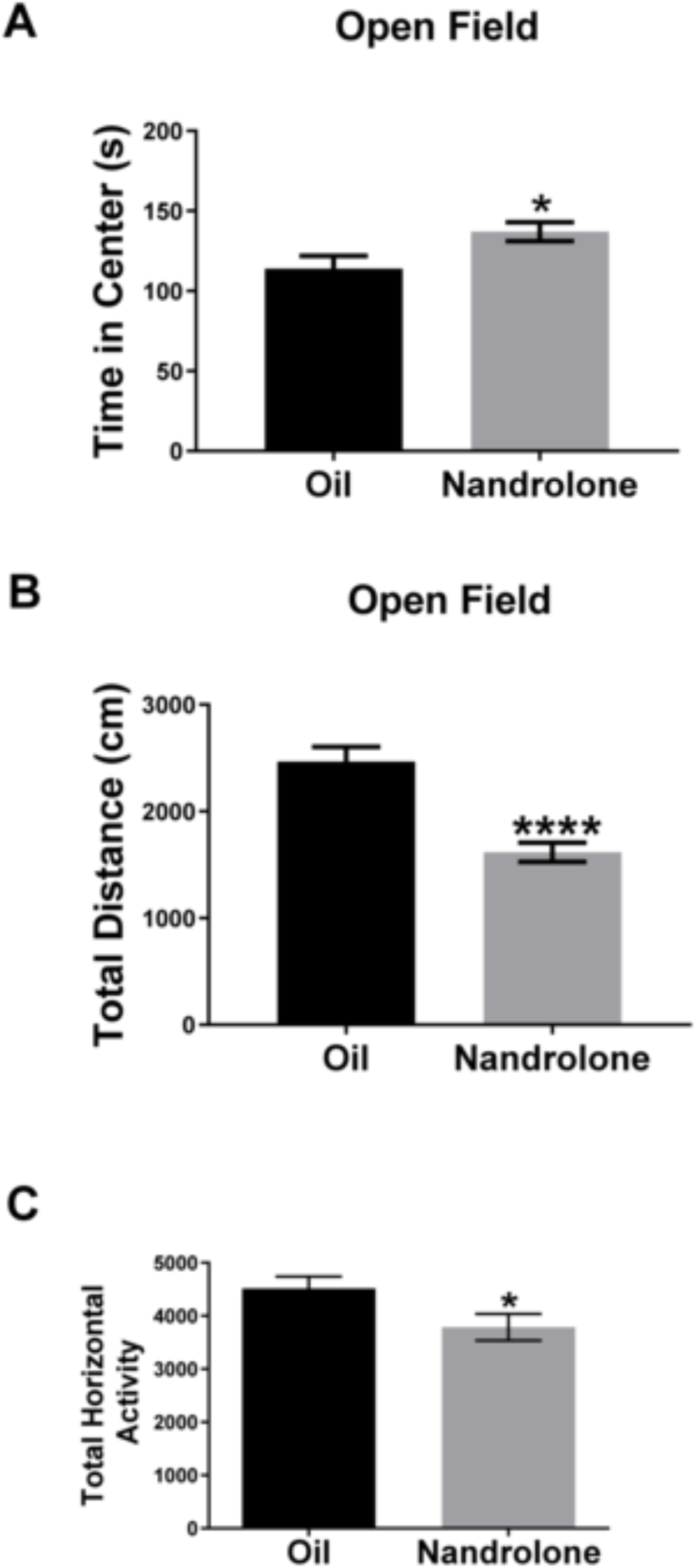
Open field activity of prepubertal rats exposed to nandrolone. Prepubertal males were injected daily with nandrolone (20 mg/kg) or sesame oil from PN 28-37. On days 0 and 1 (PN 39 and PN 40; Fig 1A), rats were tested for open field behavior. Rats were placed in the activity cage and the time spent in the center of the field, as well as the distance travelled, was recorded for 10 min using the Accuscan Versamax monitoring system. Rats treated with nandrolone (20 mg/kg) spent more time in the center of the open field (Fig 2A) and travelled less distance on days 0 (Fig 2B) and 1 (Fig 2C) than oil treated rats. Data are presented as mean ± SEM (n=10) and were analyzed with a Student’s t-Test, asterisks represent a significant difference compared to Oil group. See Table 1 for detailed statistical analysis.

**Table 1.**
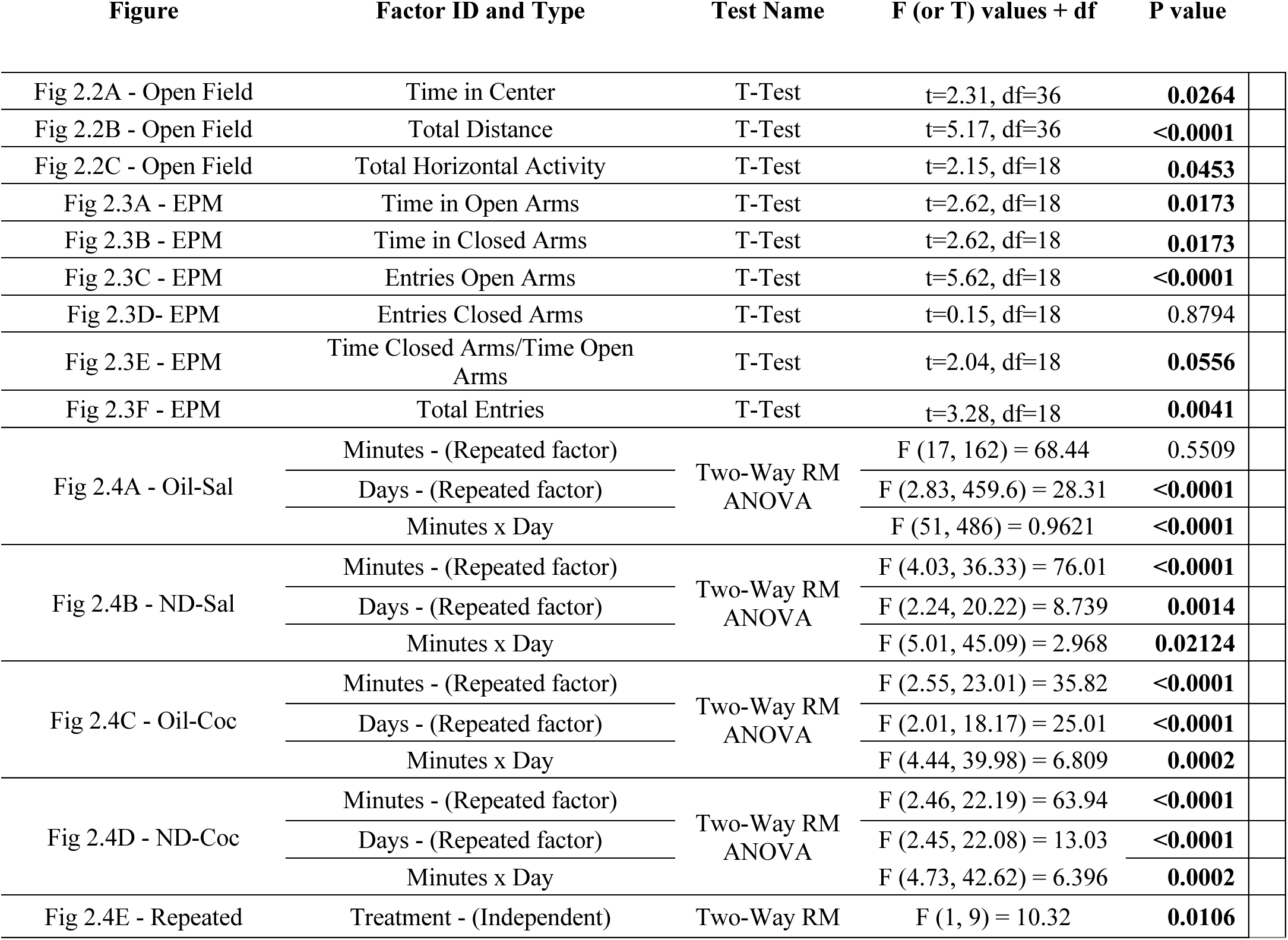
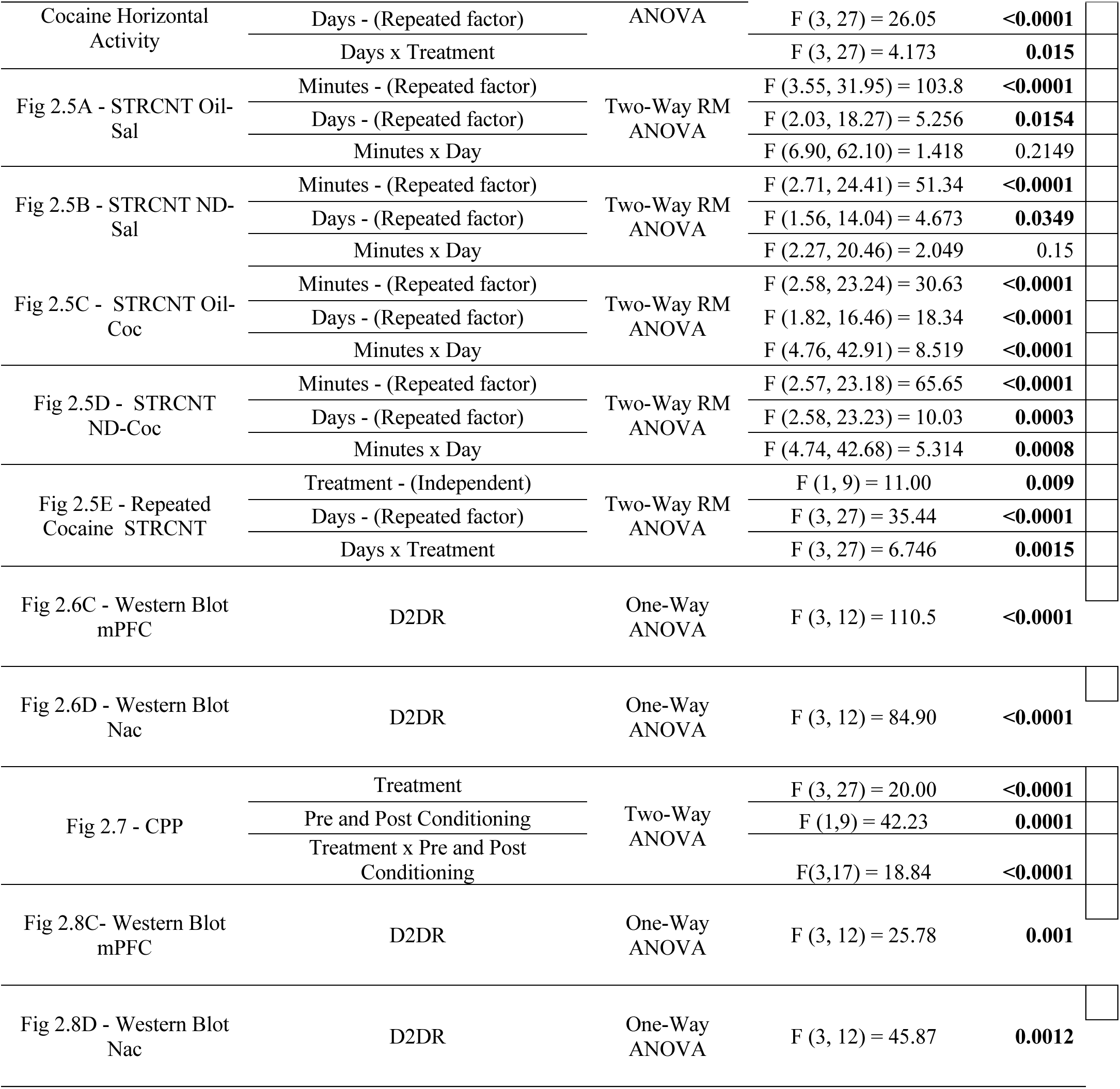
Statistical values for data analyzed by Student *t* test and ANOVA.

| Figure | Factor ID and Type | Test Name | F (or T) values + df | P value |
| --- | --- | --- | --- | --- |
| Fig 2.2A - Open Field | Time in Center | T-Test | t=2.31, df=36 | **0.0264** |
| Fig 2.2B - Open Field | Total Distance | T-Test | t=5.17, df=36 | **<0.0001** |
| Fig 2.2C - Open Field | Total Horizontal Activity | T-Test | t=2.15, df=18 | **0.0453** |
| Fig 2.3A - EPM | Time in Open Arms | T-Test | t=2.62, df=18 | **0.0173** |
| Fig 2.3B - EPM | Time in Closed Arms | T-Test | t=2.62, df=18 | **0.0173** |
| Fig 2.3C - EPM | Entries Open Arms | T-Test | t=5.62, df=18 | **<0.0001** |
| Fig 2.3D- EPM | Entries Closed Arms | T-Test | t=0.15, df=18 | 0.8794 |
| Fig 2.3E - EPM | Time Closed Arms/Time Open Arms | T-Test | t=2.04, df=18 | **0.0556** |
| Fig 2.3F - EPM | Total Entries | T-Test | t=3.28, df=18 | **0.0041** |
| Fig 2.4A - Oil-Sal | Minutes - (Repeated factor) | Two-Way RM ANOVA | F (17, 162) = 68.44 | 0.5509 |
|  | Days - (Repeated factor) |  | F (2.83, 459.6) = 28.31 | **<0.0001** |
|  | Minutes x Day |  | F (51, 486) = 0.9621 | **<0.0001** |
| Fig 2.4B - ND-Sal | Minutes - (Repeated factor) | Two-Way RM ANOVA | F (4.03, 36.33) = 76.01 | **<0.0001** |
|  | Days - (Repeated factor) |  | F (2.24, 20.22) = 8.739 | **0.0014** |
|  | Minutes x Day |  | F (5.01, 45.09) = 2.968 | **0.02124** |
| Fig 2.4C - Oil-Coc | Minutes - (Repeated factor) | Two-Way RM ANOVA | F (2.55, 23.01) = 35.82 | **<0.0001** |
|  | Days - (Repeated factor) |  | F (2.01, 18.17) = 25.01 | **<0.0001** |
|  | Minutes x Day |  | F (4.44, 39.98) = 6.809 | **0.0002** |
| Fig 2.4D - ND-Coc | Minutes - (Repeated factor) | Two-Way RM ANOVA | F (2.46, 22.19) = 63.94 | **<0.0001** |
|  | Days - (Repeated factor) |  | F (2.45, 22.08) = 13.03 | **<0.0001** |
|  | Minutes x Day |  | F (4.73, 42.62) = 6.396 | **0.0002** |
| Fig 2.4E - Repeated | Treatment - (Independent) | Two-Way RM | F (1, 9) = 10.32 | **0.0106** |
| Cocaine Horizontal Activity | Days - (Repeated factor) | ANOVA | F (3, 27) = 26.05 | **<0.0001** |
|  | Days x Treatment |  | F (3, 27) = 4.173 | **0.015** |
| Fig 2.5A - STRCNT Oil-Sal | Minutes - (Repeated factor) | Two-Way RM ANOVA | F (3.55, 31.95) = 103.8 | **<0.0001** |
|  | Days - (Repeated factor) |  | F (2.03, 18.27) = 5.256 | **0.0154** |
|  | Minutes x Day |  | F (6.90, 62.10) = 1.418 | 0.2149 |
| Fig 2.5B - STRCNT ND-Sal | Minutes - (Repeated factor) | Two-Way RM ANOVA | F (2.71, 24.41) = 51.34 | **<0.0001** |
|  | Days - (Repeated factor) |  | F (1.56, 14.04) = 4.673 | **0.0349** |
|  | Minutes x Day |  | F (2.27, 20.46) = 2.049 | 0.15 |
| Fig 2.5C - STRCNT Oil-Coc | Minutes - (Repeated factor) | Two-Way RM ANOVA | F (2.58, 23.24) = 30.63 | **<0.0001** |
|  | Days - (Repeated factor) |  | F (1.82, 16.46) = 18.34 | **<0.0001** |
|  | Minutes x Day |  | F (4.76, 42.91) = 8.519 | **<0.0001** |
| Fig 2.5D - STRCNT ND-Coc | Minutes - (Repeated factor) | Two-Way RM ANOVA | F (2.57, 23.18) = 65.65 | **<0.0001** |
|  | Days - (Repeated factor) |  | F (2.58, 23.23) = 10.03 | **0.0003** |
|  | Minutes x Day |  | F (4.74, 42.68) = 5.314 | **0.0008** |
| Fig 2.5E - Repeated Cocaine STRCNT | Treatment - (Independent) | Two-Way RM ANOVA | F (1, 9) = 11.00 | **0.009** |
|  | Days - (Repeated factor) |  | F (3, 27) = 35.44 | **<0.0001** |
|  | Days x Treatment |  | F (3, 27) = 6.746 | **0.0015** |
| Fig 2.6C - Western Blot mPFC | D2DR | One-Way ANOVA | F (3, 12) = 110.5 | **<0.0001** |
| Fig 2.6D - Western Blot Nac | D2DR | One-Way ANOVA | F (3, 12) = 84.90 | **<0.0001** |
| Fig 2.7 - CPP | Treatment | Two-Way ANOVA | F (3, 27) = 20.00 | **<0.0001** |
|  | Pre and Post Conditioning |  | F (1,9) = 42.23 | **0.0001** |
|  | Treatment x Pre and Post Conditioning |  | F(3,17) = 18.84 | **<0.0001** |
| Fig 2.8C- Western Blot mPFC | D2DR | One-Way ANOVA | F (3, 12) = 25.78 | **0.001** |
| Fig 2.8D - Western Blot Nac | D2DR | One-Way ANOVA | F (3, 12) = 45.87 | **0.0012** |

### 4.2 Elevated Plus Maze

Nandrolone administration decreased the time spent (Fig 3A: Student T-test: t=2.623, df=18, p = 0.0173) into the closed arms of the EPM compared to rats that received oil. In addition, nandrolone decreased the total number of arm entries (Fig 3B: Student’s T-test: t=3.28, df=18, p=0.0041), which is considered an indication of decreased locomotor activity.

**Figure 3.**
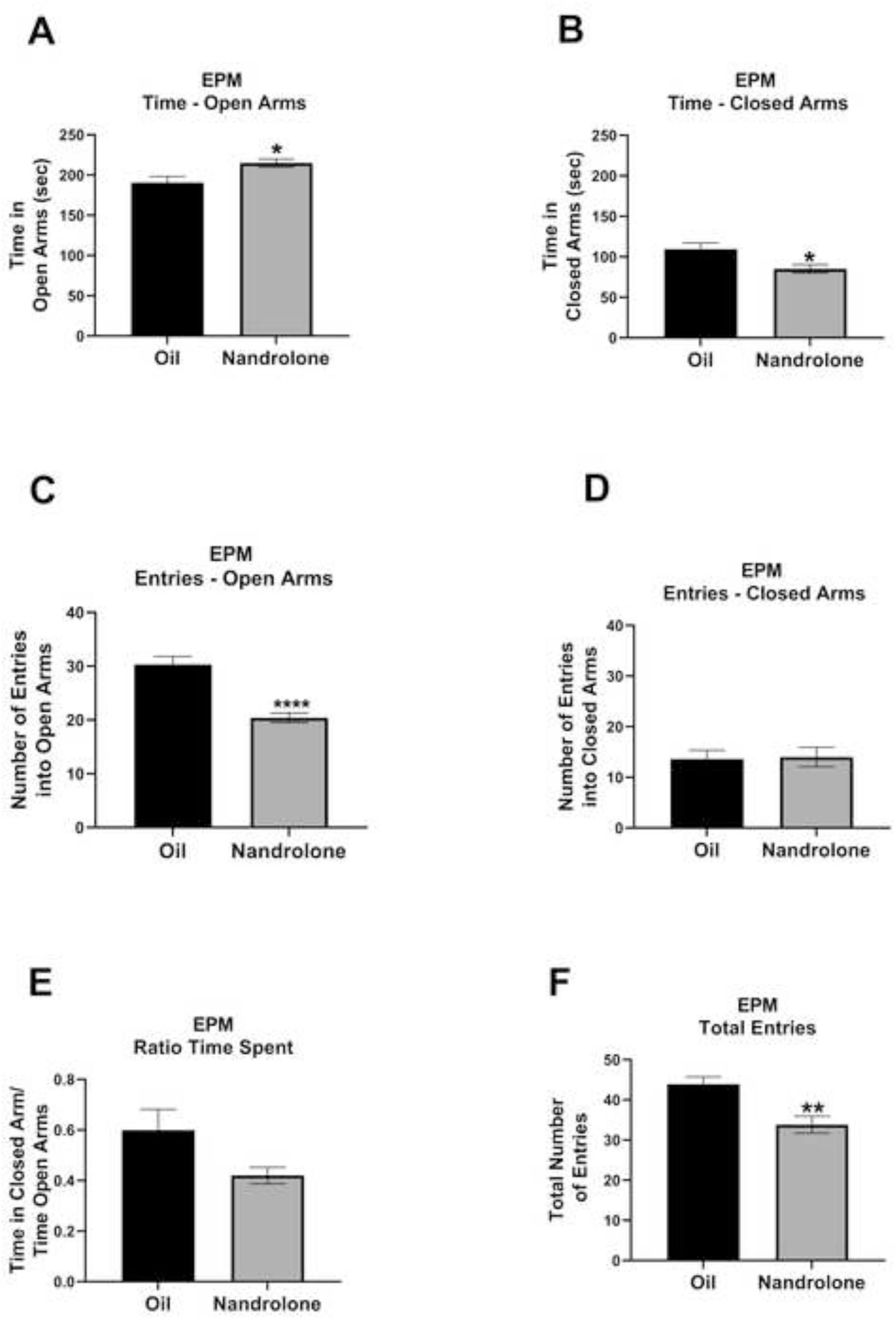
Results from the EPM test of prepubertal rats treated with nandrolone. Male rats were injected daily with nandrolone (20 mg/kg) or sesame oil from PN 28-37 and tested in the elevated plus maze on day 38. Rats were placed at the junction between the open and closed arms and behavioral activity was recorded for 5 min using the ANY-Maze™ tracking software. Rats treated with nandrolone spent less time in the closed arms (Fig 3A), an indication of decreased anxiety. In addition, rats that received nandrolone had fewer total entries into both arms than oil-treated rats (Fig 3B), an indication of reduced ambulation. Data are presented as mean ± SEM (n=10) and analyzed with a Student t-Test. Asterisks represent a significant difference compared to Oil treated rats.

### 4.3 Behavioral sensitization

Rats treated with nandrolone showed a higher locomotor response to cocaine on day 5 than that displayed on day 1, i.e. they displayed behavioral sensitization (Fig 4A, 4B, 4C, 4D, 4E). Of the 12 cocaine-induced locomotor activity timepoints measured during the 60 min after injection (5 min intervals), 8 were higher in rats that received nandrolone-cocaine (Fig 4C vs 4D). This pattern was maintained on day 13, 10 out of 12 cocaine-induced locomotor activity timepoints for nandrolone-cocaine treated rats were higher from those on day 1 (Fig 4D). In comparison, oil-cocaine treated rats did not show differences in cocaine-induced locomotor activity between days 1 and 5 in any of the 12 timepoints measured and showed differences in only 5 of the 12 timepoints comparing day 1 with day 13 (Fig 4C). Thus, rats treated with nandrolone-cocaine displayed behavioral sensitization earlier (day 5) than oil-cocaine treated rats (day 13) (Fig.4D: Two-Way RM ANOVA, Days, F_(2.45, 22.08)_ = 13.03; p<0.0001). Similar results were obtained with Stereotyped activity (Fig.5A-E: Two-Way RM ANOVA, Days, F_(2.58, 23.23)_ = 10.03, p= 0.0003). In addition, cocaine-induced locomotor activity of nandrolone-treated rats was greater than that of oil-treated rats in 9 of the 12 timepoints measured on day 5.

**Figure 4.**
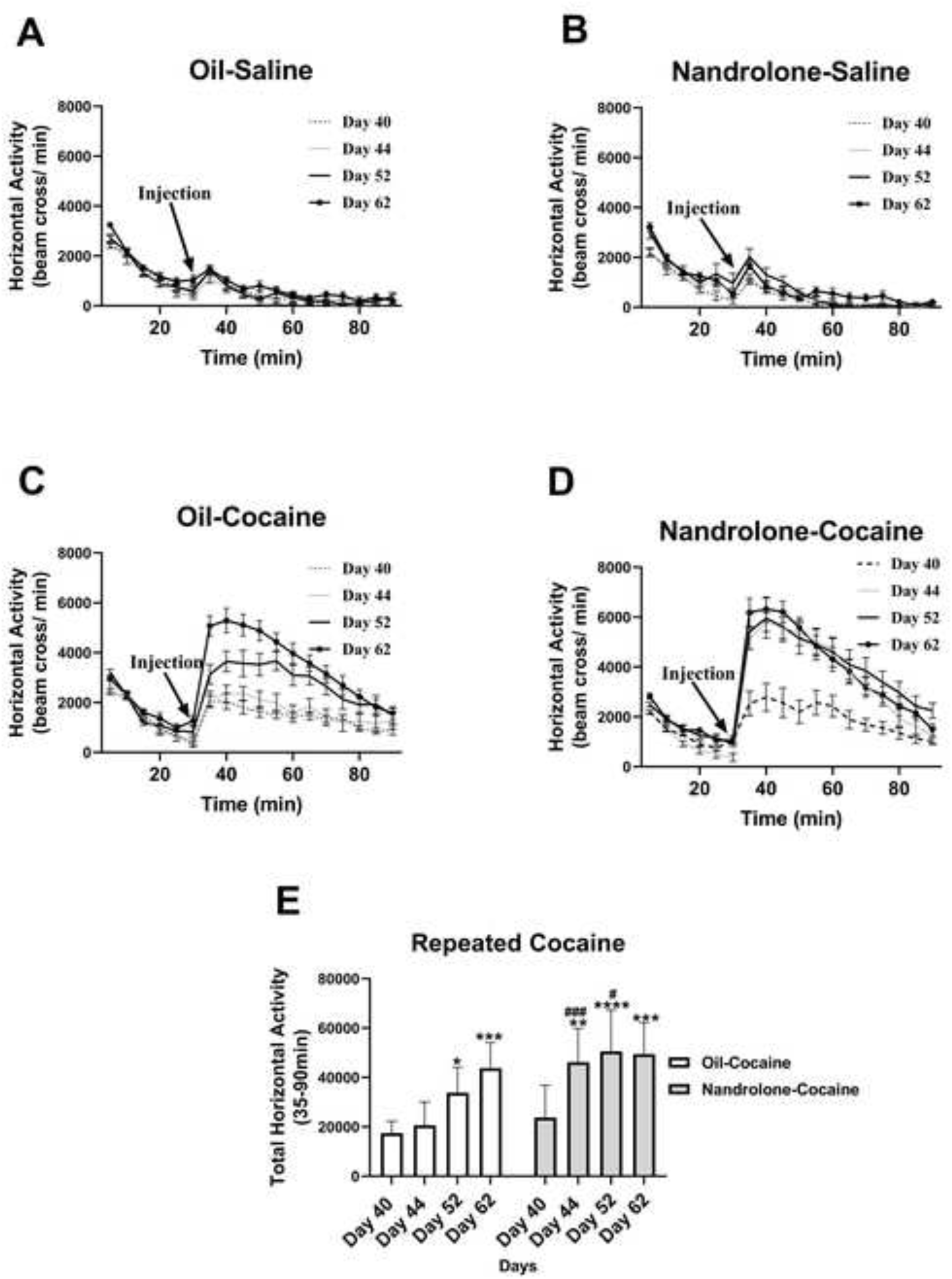
Cocaine-induced locomotor activity of rats exposed to nandrolone during postnatal days 28-37. Cocaine-induced locomotor activity of prepubertal rats injected daily from PN 28-37 with nandrolone (20 mg/kg) or sesame oil and tested for behavioral sensitization to cocaine from PN 40-62. **Figs 4A and 4B**. Timecourse of locomotor activity of saline (Fig 4A) and nandrolone (Fig 4B) treated males. No differences between oil and nandrolone-treated males were observed. **Fig 4C**. A significant increase in cocaine-induced locomotor activity was observed when comparing the timecourses of day 40 vs day 52 and day 40 vs day 62 in Oil-treated males. Thus, these males displayed behavioral sensitization by day 52. **Fig 4D**. A significant increase in cocaine-induced locomotor activity was observed when comparing the timecourses of day 40 vs day 44, vs day 52 and vs day 62 in nandrolone-treated males. These results suggest that nandrolone accelerates the maturation of the neural circuitry that regulates behavioral sensitization. Data are presented as mean ± SEM and analyzed with a Repeated Measures ANOVA using Tukey’s multiple comparison for post-hoc analysis. (See Supplemental Tables 1, 2, and 4 for complete statistical analysis values). **Fig 4E**. Rats that received nandrolone show a robust increase in cumulative cocaine-induced locomotor activity by day 44, an increase that is maintained after two withdrawal periods. In contrast, it is at day 52 that oil-treated rats show a difference that is further increased by day 62. Data are presented as mean ± SEM (n=10). Repeated Measures Two-Way ANOVA: F_(3,27)_ = 4.173, p = 0.0150. **Oil-Coc**: Day 40 vs Day 44, p = 0.9968; Day 40 vs Day 52, p = 0.0288; Day 40 vs Day 62, p = 0.0001. **ND-Coc**: Day 40 vs Day 44, p = 0.0011; Day 40 vs Day 52, p = <0.0001; Day 40 vs Day 62 p = 0.0002. **Oil-Coc vs ND-Coc**: Day 40, p = 0.8684; Day 44, p = 0.0002; Day 52, p = 0.0230; Day 62 p = 0.9209. P < 0.05 was considered statistically significant Day 40 vs. Day 44, Day 52 and Day 62. (*): significantly different within groups; (#): significantly different between groups (Oil-Cocaine vs Nandrolone-Cocaine).

**Figure 5.**
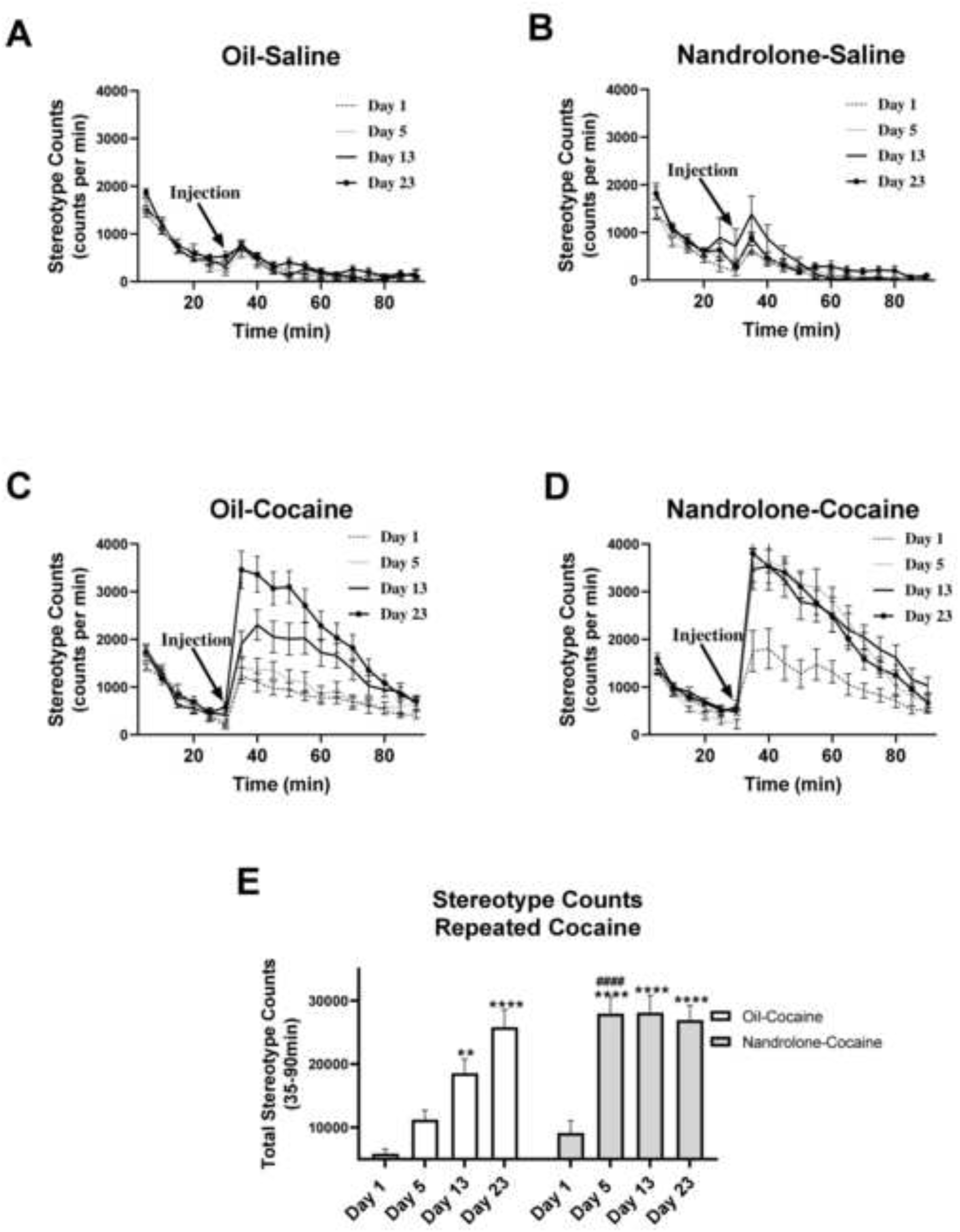
Cocaine-induced stereotyped activity of rats exposed to nandrolone during postnatal days 28-37. Cocaine-induced stereotyped activity of prepubertal rats injected daily from PN 28-37 with nandrolone (20 mg/kg) or sesame oil and tested for behavioral sensitization to cocaine from PN 40-62. **Figs 5A and 5B**. Timecourse of stereotyped activity of saline (Fig 5A) and nandrolone (Fig 5B) treated males. No differences between oil and nandrolone-treated males were observed. **Fig 5C**. A significant increase in cocaine-induced stereotyped activity was observed when comparing the timecourses of day 40 vs day 52 and day 40 vs day 62 in Oil-treated males. Thus, saline males displayed behavioral sensitization by day 52. **Fig 5D**. A significant increase in cocaine-induced locomotor activity was observed when comparing the timecourses of day 40 vs day 44, vs day 52 and vs day 62 in nandrolone-treated males. Similar to what we observed with total horizontal activity, these results indicate that nandrolone accelerates the development of behavioral sensitization. Data are presented as mean ± SEM (n=10) and analyzed with a Repeated Measures Two-Way ANOVA using Tukey’s multiple comparison for post-hoc analysis. See Tables 1, 2, and 3 for complete statistical analysis values. **Fig 5E**. Rats that received nandrolone show a robust increase in cumulative cocaine-induced stereotyped activity by day 44, an increase that is maintained after two withdrawal periods. In contrast, it is at day 52 that oil-treated rats show a difference that is further increased by day 62. Data are presented as mean ± SEM. Repeated Measures Two-Way ANOVA: F_(3,27)_ = 6.746, p = 0.0015. **Oil-Coc**: Day 40 vs Day 44, p = 0.5042; Day 40 vs Day 52, p = 0.015; Day 40 vs Day 62, p < 0.0001. **ND-Coc**: Day 40 vs Day 44, p < 0.0001; Day 40 vs 52, p = <0.0001; Day 40 vs Day 62, p < 0.0002. **Oil-Coc vs ND-Coc**: Day 40, p = 0.9224; Day 44, p < 0.0001; Day 52, p = 0.0281; Day 62 p = 0.9999. P < 0.05 was considered statistically significant Day 40 vs. Day 44, Day 52 and Day 62. (*): significantly different within groups; (#): significantly different between groups (Oil-Cocaine vs Nandrolone-Cocaine).

**Table 2.**
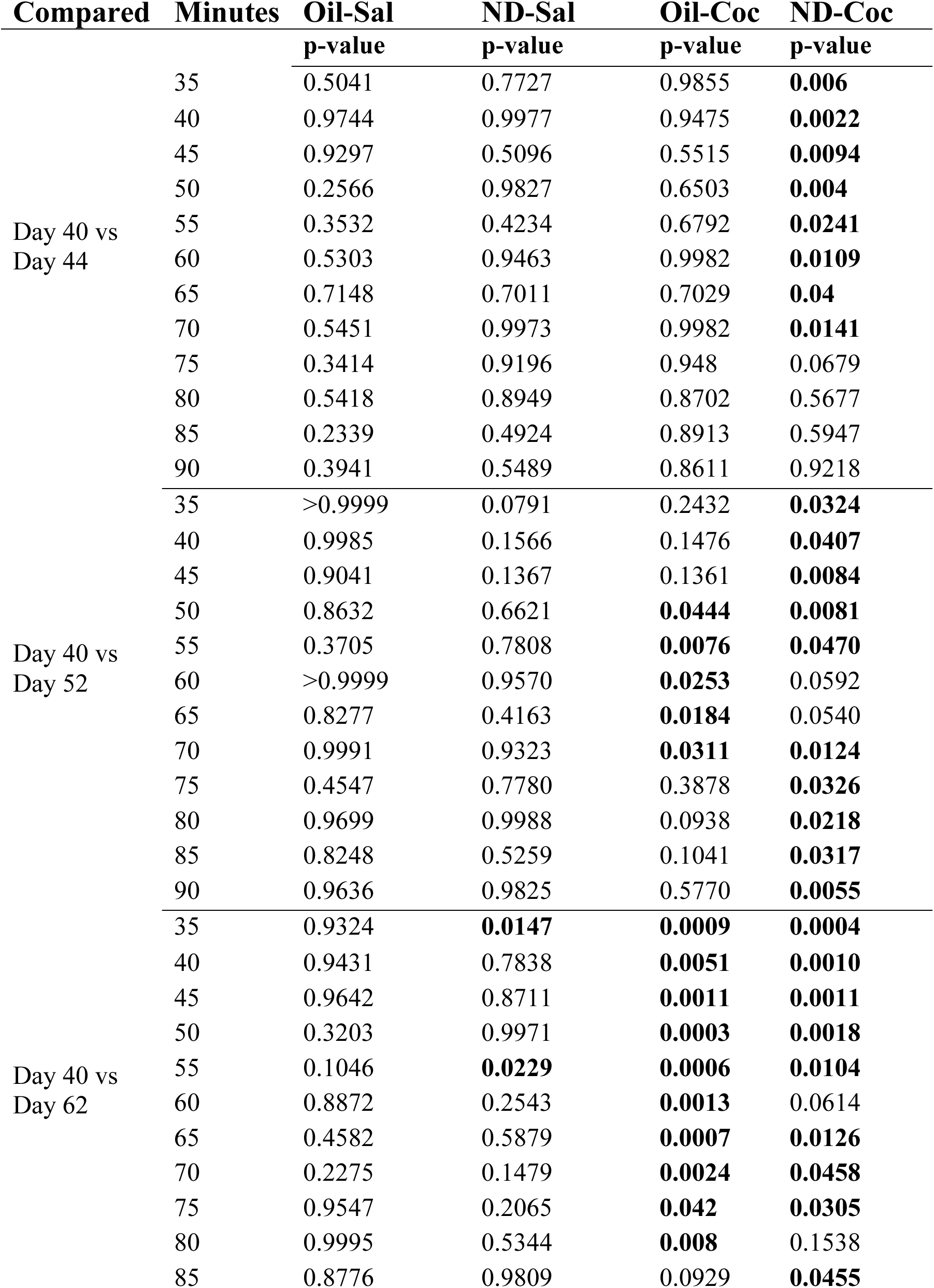
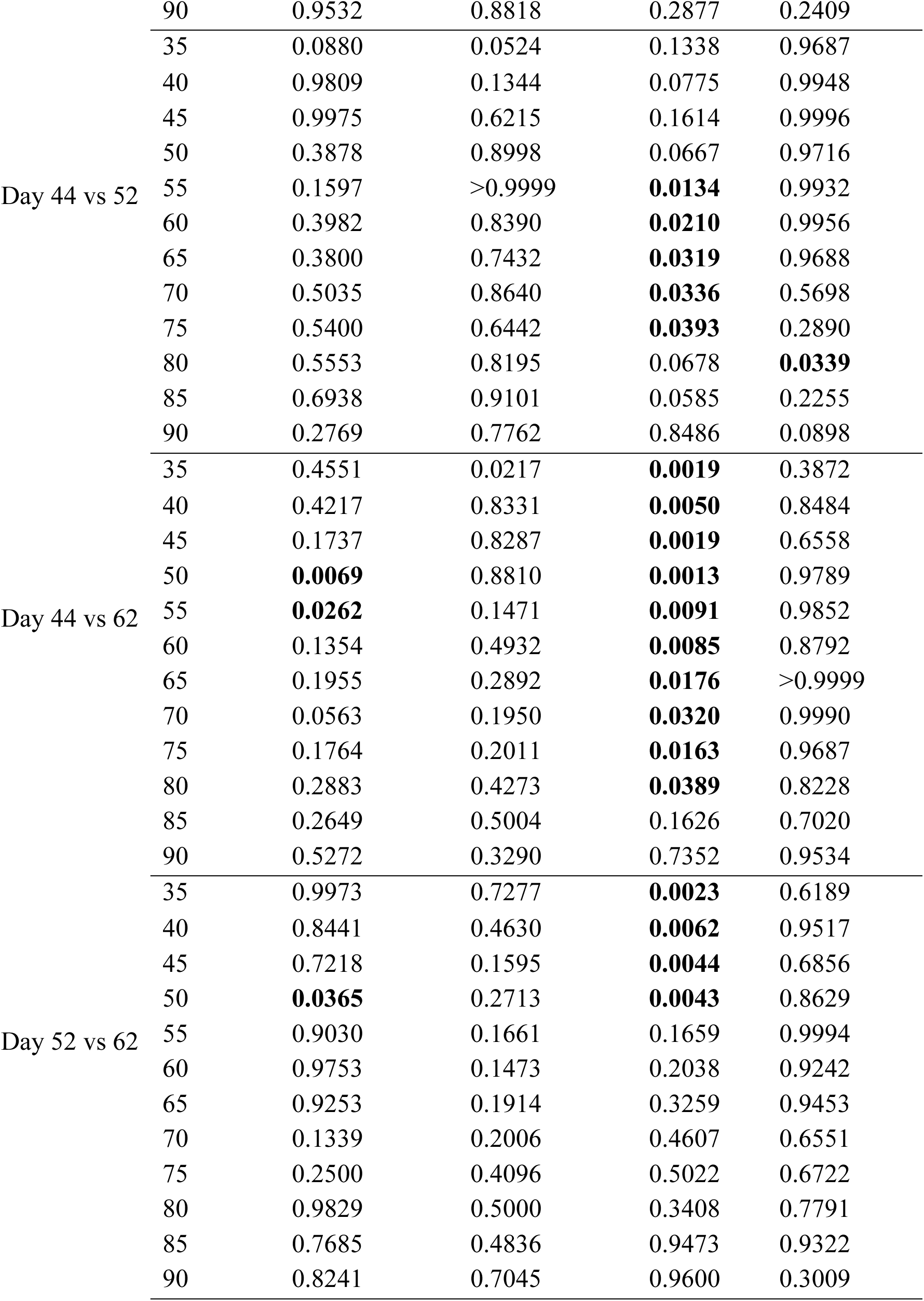
Two-Way ANOVA Post-hoc Analysis of Horizontal Activity Days.

**Table 3.**
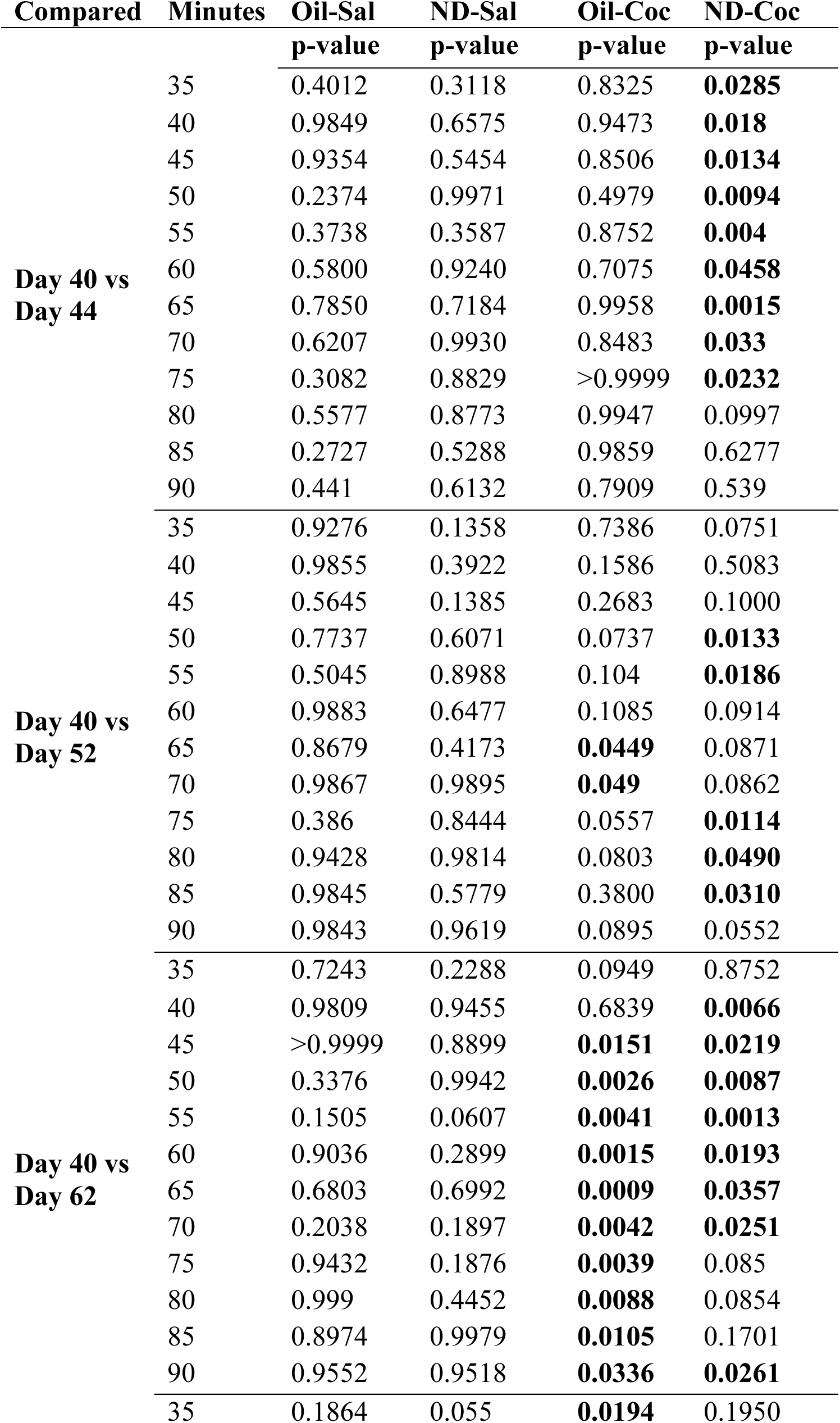
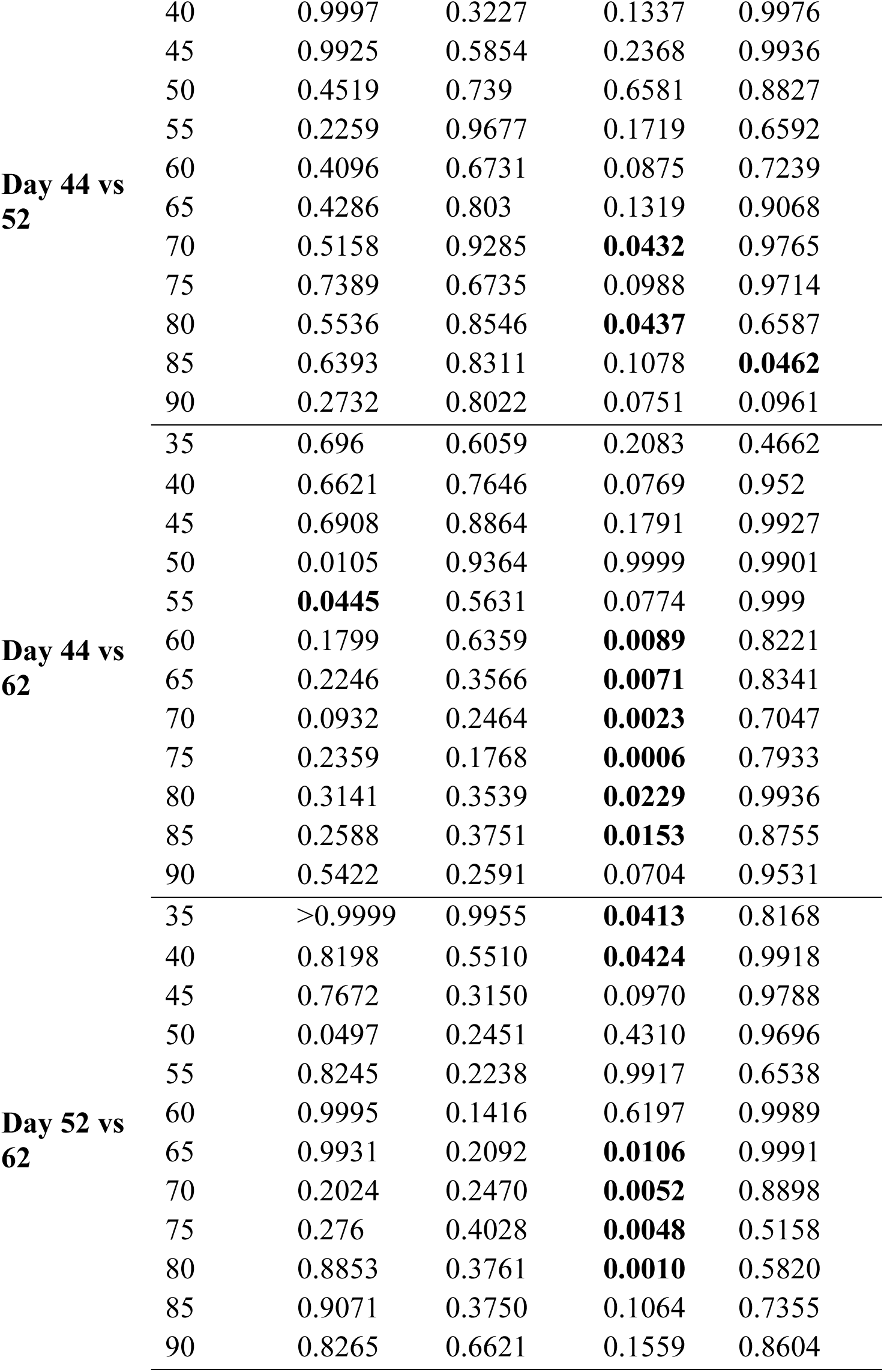
Two-Way ANOVA Post-hoc Analysis of Stereotyped Counts Days.

### 4.4 Western Blot Analysis of D2DR in mPFC (Sensitization brains)

Our data show that cocaine decreased D2DR in the mPFC (Fig 6A and Fig 6C). This effect was not altered by preexposure to nandrolone (Fig. 6A and 6C): One Way ANOVA, Tukey’s multiple comparisons, F _(3, 12)_ = 110.5, <0.0001; Oil-Sal vs ND-Sal, p = 0.1559; Oil-Sal vs Oil-Coc, p < 0.0001; Oil-Sal vs ND-Coc, p < 0.0001; ND-Sal vs Oil-Coc, p < 0.0001; ND-Sal vs ND-Coc, p < 0.0001; Oil-Coc vs ND-Coc, p = 0.9909.

**Figure 6.**
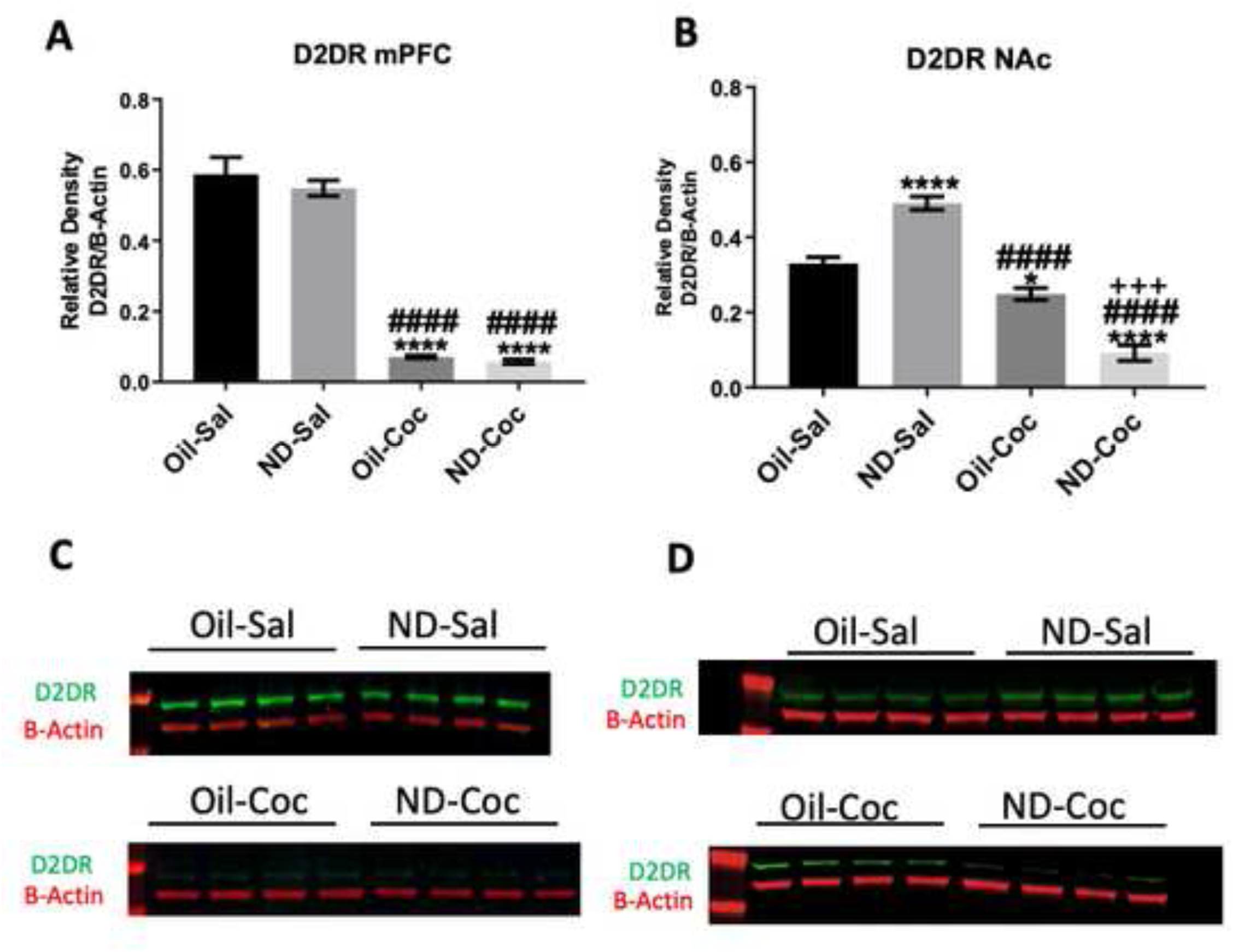
D2DR in the mPFC and NAc of rats exposed to nandrolone during postnatal days 28-37 and tested for behavioral sensitization. **Fig. 6A and Fig 6C**: Nandrolone did not affect D2DR expression in the mPFC, however a decrease in D2DR expression in the mPFC was observed in animals treated with cocaine. **Fig. 6B and Fig 6D**: In contrast, nandrolone treatment increased D2DR expression in the NAc and similar to what was observed in the PFC, cocaine reduced D2DR in the NAc. Data are presented as mean ± SEM (n=4). Representative western blots of D2DR and B-Actin in the mPFC **(Fig 6C)** and NAc **(Fig 6D)** of rats exposed to nandrolone during postnatal days 28-37, tested for behavioral sensitization and sacrificed on PN 64. Data were analyzed using One-Way ANOVA. (See Supplemental Table 1 for statistical analysis).

### 4.5 Western Blot Analysis of D2DR in NAc (Sensitization brains)

Rats treated with nandrolone showed increased expression of D2DR in the NAc (Fig 6B and 6D). Cocaine treatment decreased the expression of D2DR in this brain area; this decrease was more pronounced in animals that received nandrolone (Fig.6B and 6D: One Way ANOVA, Tukey’s multiple comparisons, F(3, 12) = 84.90, < 0.0001; Oil-Sal vs ND-Sal, p = 0.0002; Oil-Sal vs Oil-Coc, p = 0.0339; Oil-Sal vs ND-Coc, p < 0.0001; ND-Sal vs Oil-Coc, p < 0.0001; ND-Sal vs ND-Coc, p < 0.0001; Oil-Coc vs ND-Coc, p = 0.0002.

### 4.6 Conditioned Place Preference

All animals that were tested displayed CPP to cocaine, however nandrolone decreased the time spent in the chamber associated with cocaine (Fig 7). Rats treated with oil-cocaine displayed CPP to cocaine more robustly than nandrolone-cocaine treated rats (Fig.7). During postconditioning, oil treated rats spent 70% of their time in the side associated with cocaine vs 31% during the preconditioning phase. In contrast, rats treated with nandrolone spent 48% of their time during postconditioning in the chamber associated with the drug vs 30% during preconditioning. Data were analyzed with a Two-Way ANOVA, with pre and post-conditioning as repeated measures = F_(1, 62)_= 28.22, p <0.0001 and Cocaine as the independent factor. Cocaine’s effect: F_(3,62)_ = 21.97, p < 0.0001; Oil-Coc pre vs post conditioning, p < 0.0001; ND-Coc pre vs post conditioning, p = 0.0397; Oil-Coc post vs ND-Coc post conditioning, p = 0.0035.

**Figure 7.**
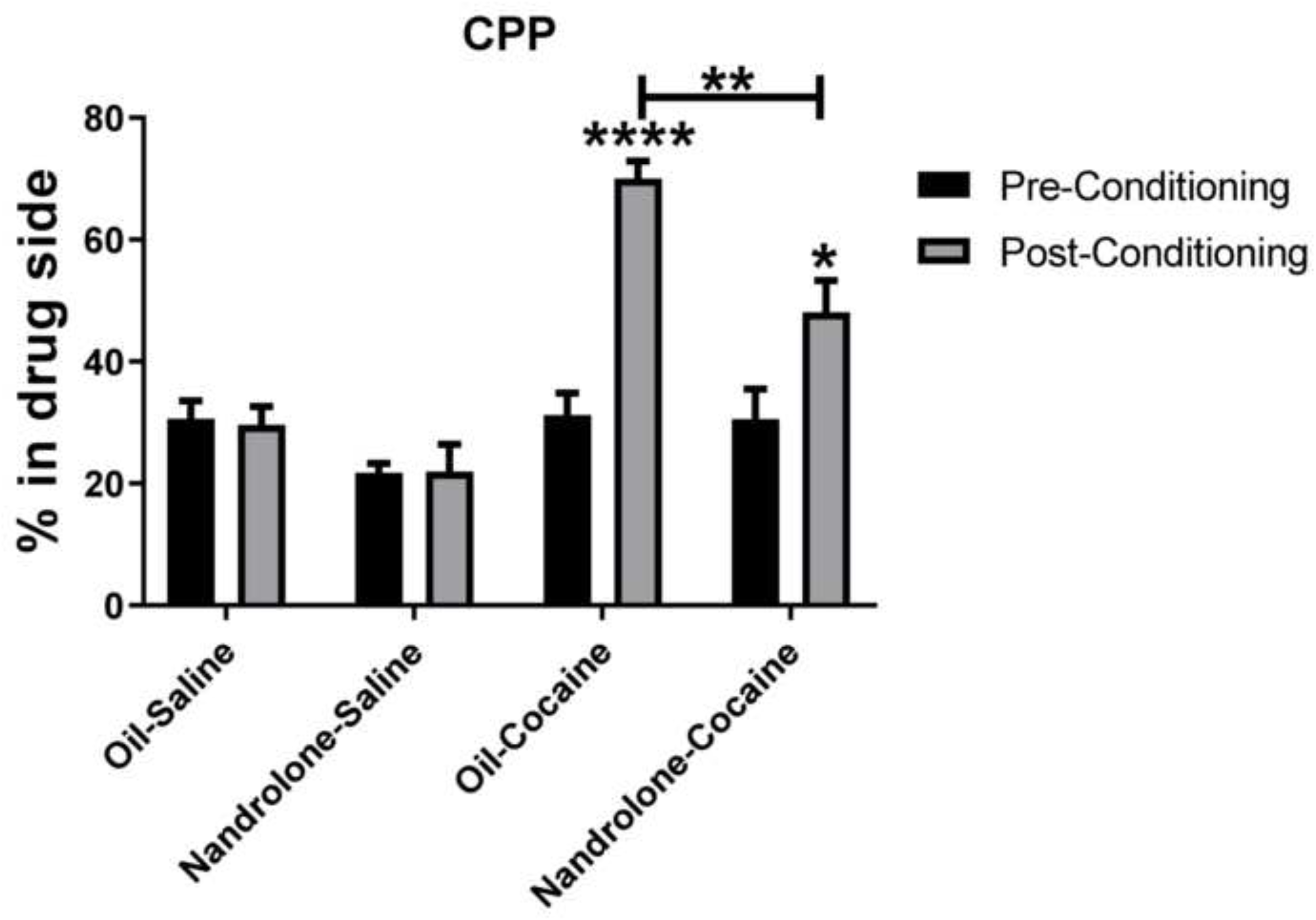
Conditioned Place Preference to cocaine of rats exposed to nandrolone during postnatal days 28-37. Adolescent male rats were injected daily from PN 28-37 with nandrolone (20 mg/kg) or sesame oil and tested for CPP to cocaine from PN 40-53. Nandrolone-treated males showed a decrease in the time spent in the chamber associated with cocaine compared to oil-treated males. During the postconditioning test, nandrolone-treated rats spent 48% of their time in the chamber associated with cocaine compared to 70% spent by oil-treated males. Rats injected with saline did not show a change in the time spent in the chamber where they were injected with saline. Although oil and nandrolone treated males conditioned to cocaine, conditioning was more robust in Oil-treated males. Data are presented as mean ± SEM (n=8) and analyzed by a Two Way ANOVA. See Supplemental Table 1 for statistical analysis details.

### 4.7 Western Blot Analysis for D2DR in mPFC (CPP brains)

Similar to our previous results, rats that received cocaine had decreased D2DR in the PFC compared to saline treated rats (Fig 8A y 8C). However, the brains from nandrolone-treated males used in the CPP experiments had more D2DR in the PFC than oil-treated rats (Fig 8A y 8C). The difference from these two experiments is that the animals used for the CPP experiments were killed at day 53 and those used for the sensitization experiments were sacrificed 11 days later, at day 64. In addition, for the sensitization studies rats received 7 cocaine injections (days 1-5, 13 and 23), whereas for the CPP experiments rats received 5 cocaine injections (days 2,4,6,8,10). Fig. 8C: One Way ANOVA, Tukey’s multiple comparisons, F_(3, 12)_ = 28.78, <0.0001; Oil-Sal vs ND-Sal, p = 0.1752; Oil-Sal vs Oil-Coc, p = 0.0013; Oil-Sal vs ND-Coc, p = 0.0007; ND-Sal vs Oil-Coc, p < 0.0001; ND-Sal vs ND-Coc, p < 0.0001; Oil-Coc vs ND-Coc, p = 0.9825.

**Figure 8.**
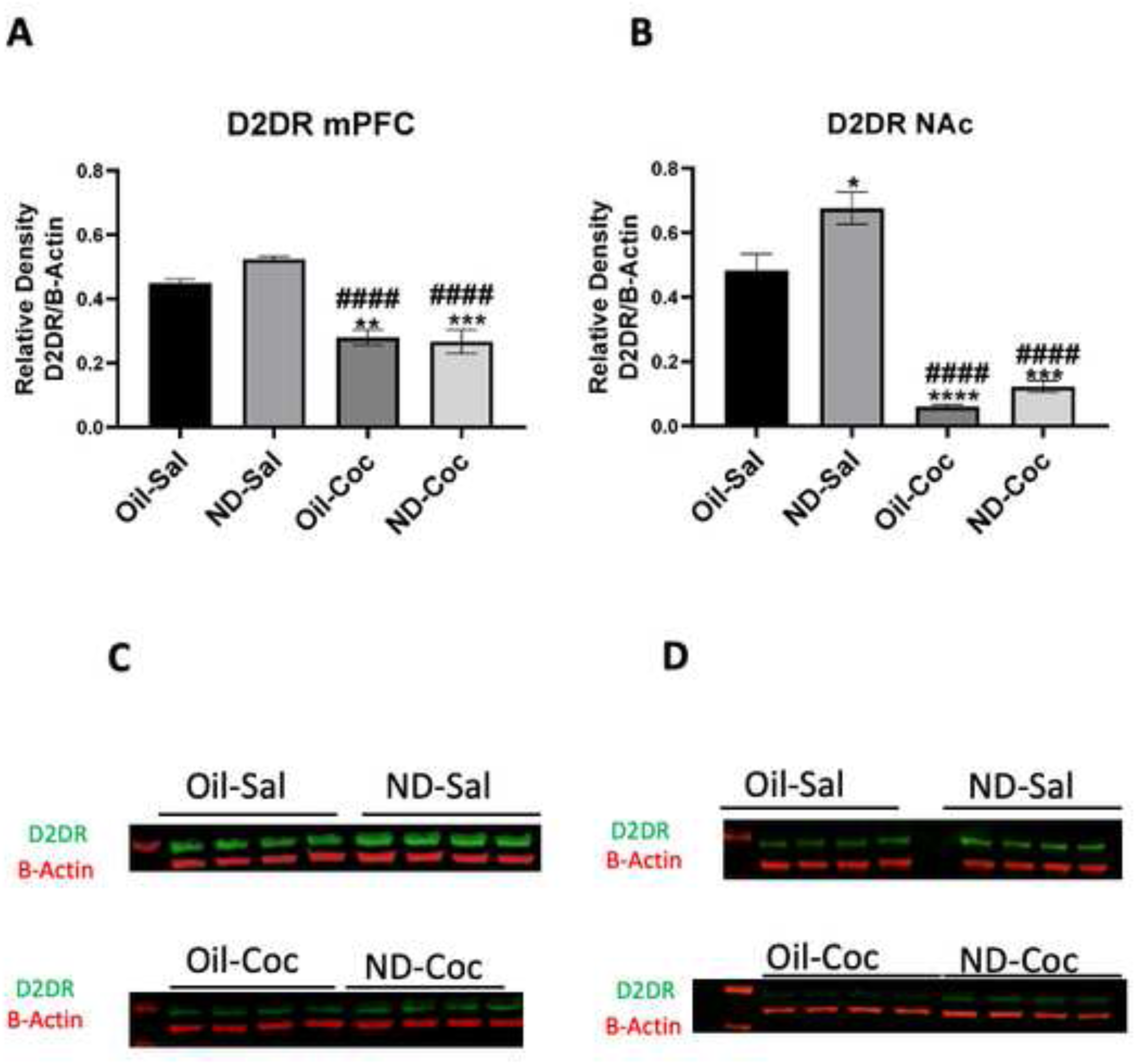
D2DR in the mPFC and NAc of rats exposed to nandrolone during postnatal days 28-37 and tested for CPP. **Fig. 8B and 8D**: Nandrolone increased D2DR expression in the mPFC and NAc of prepubertal rats. **Fig. 8B and Fig 8D**: In contrast, cocaine decreased D2DR expression in both brain areas (mPFC and NAc), similar to what was observed in brains obtained from animals that were tested for behavioral sensitization and killed at a later age (PN 64). Representative western blots of D2DR and B-Actin in the mPFC **(Fig 8C)** and NAc **(Fig 8D)** of rats exposed to nandrolone during postnatal days 28-37, tested for CPP and sacrificed on PN 53. Data are presented as mean ± SEM (n=4) and analyzed using One-Way ANOVA. (See Supplemental Table 1 for detailed statistical analysis).

### 4.8 Western Blot Analysis for D2DR in NAc (CPP brains)

Similar to the results obtained from the brains of animals sensitized to cocaine (Fig 6D) CPP rats treated with nandrolone showed increased expression of D2DR in the NAc. In contrast, cocaine treatment decreased the expression of D2DR in this brain area, similar to our previous results. Fig.8D: One Way ANOVA, Tukey’s multiple comparisons, F (3, 12) = 45.87, <0.0001; Oil-Sal vs ND-Sal, p =0.0316; Oil-Sal vs Oil-Coc, p < 0.0001; Oil-Sal vs ND-Coc, p = 0.0001; ND-Sal vs Oil-Coc, p < 0.0001; ND-Sal vs ND-Coc, p < 0.0001; Oil-Coc vs ND-Coc, p = 0.6828.

## 5. Discussion

### 5.1 Summary of results

Nandrolone was administered prior to puberty (PN 28-37) and the effects on anxiety and risk-taking behaviors (days 38 and 39), and on the response to cocaine (day 40) was assessed immediately after nandrolone treatment, not after a withdrawal period. We found that administration of nandrolone decanoate decreased anxiety and altered addictive-like behaviors in male rats.

Prepubertal (PN 28-37) males treated with nandrolone showed reduced anxiety and basal locomotor activity compared to oil-treated males. In contrast, nandrolone increased risk-taking behavior and the locomotor response to cocaine on day 5 and accelerated the development of cocaine sensitization in young animals (i.e., they required less cocaine injections). Nandrolone also attenuated CPP to cocaine. Changes in D2DR in the NAc may partially mediate these behavioral changes: an increase in D2DR in the NAc was observed in nandrolone-treated males, whereas those treated with cocaine or with nandrolone and cocaine had less D2DR in the NAc and PFC.

### 5.2 Anxiety and risk-behaviors

Previous studies have found that the use of AAS is associated with adverse effects on emotion, cognition, and rewarding behaviors (Grönbladh et al., 2016). Clinical studies indicate that most of the effects of AAS on anxiety appear after cessation of use, which may be at the end of an AAS cycle, or after longer withdrawal periods (Pope et al., 1996). People who feel depressed or are experimenting withdrawal symptoms, are more likely to revert and use anabolic steroids or other drugs of abuse (Malone et al., 1995).

Other studies indicate that 60% of children born of mothers with polycystic ovarian syndrome, which results in increased androgen exposure during fetal development, have a higher prevalence of psychiatric disorders, such as depression and anxiety (Dokras et al., 2012). Additional studies in rodents confirm that exposure to excessive androgens during prenatal development increases the number of offspring that subsequently display anxiety behaviors (Hu et al., 2015). In addition, several psychiatric disorders like anxiety, depression, mood disorders, psychosis and substance abuse tend to manifest themselves close to puberty (Paus et al., 2008)

Very few studies have examined the effects of AAS administered prepubertally on emotional behaviors, although the use of AAS in this population is very high. Our study found that nandrolone exposure prepubertally decreased anxiety-like behaviors. Rats treated with nandrolone showed a decrease in the time spent in the closed arms of an EPM and an increase in the center of an open field compared to oil-treated rats, behaviors associated with decreased anxiety.

This increase in time spent in the center of an open field is associated with increased risk-taking behavior as animals that venture more into the center of an open field are at greater risk of being detected by predators.

Previous studies such as Kouvelas et al (2008) have found that nandrolone administration (15mg/kg) to adult rats decreased anxiety as measured in the EPM. Decreased anxiety following androgen administration has also been reported by others (Aikey et al., 2002; Bitran et al., 1993; Rainer et al., 2014). This anxiety-lowering effect has also been seen with androgens such as testosterone and DHT (Bitran et al., 1993; Edinger and Frye, 2006).

However, it is not unequivocally established whether AAS have anxiolytic or anxiogenic properties since several studies in adult male rats (Rosic et al., 2014) or in adolescent rats tested as adults (Ricci et al., 2012; Morrison et al., 2015) report that AAS exert anxiogenic effects (Ricci et al., 2012; Morrison et al., 2015).

The difference between studies investigating the effect of AAS on anxiety may vary depending on the type of androgen administered, the age of exposure, the duration of treatment, and the dose administered. These last studies used mainly an aromatizable form of androgen, and although the doses used were smaller (5 mg/kg vs 20 mg/kg), as well as the total dose administered (60, 150 or 175 mg/kg vs 200 mg/kg) the duration of treatment was longer (30-35 days vs 10 days). In addition, our current study is unique in that nandrolone was administered prior to puberty (PN 28-37) and the effects on anxiety and risk-taking behaviors assessed the day after the last injection (days 38 and 39) and not after a withdrawal period.

Studies that examined the role of androgens on anxiety report that androgen receptors, especially those in hippocampal tissue, appear to play a significant role in mediating this response (Edinger and Frye, 2004). Kouvelas et al., (2008) found that the anxiolytic effect of AAS was abolished by treatment with flutamide, an androgen receptor inhibitor.

A correlation between high-risk behaviors in adolescents and testosterone has also been established. For example, a positive correlation was found between salivary testosterone and activation of the nucleus accumbens in response to a monetary reward in adolescent boys ages 10 to 16 (Op De MacKs et al., 2011). Not only do testosterone levels correlate positively with receiving a financial reward, but AAS users also engage more frequently in other risky behaviors such as gambling.

### 5.3 Locomotor activity

Rats treated with nandrolone showed a decrease in distance traveled in an open field and a reduction in the total number of arm entries when tested in an EPM, indicative of lower ambulation. Lower locomotor activity of nandrolone-treated males was also evident during the first 10 min of habituation (day 0) and on day 1 of the sensitization trial (Fig.3C). This effect of nandrolone on locomotor activity appears to be related to novelty-induced exploration since it was not observed upon further testing (days 5, 13 and 23).

Most studies agree that androgens decrease ambulation. In a similar study, prepubertal males treated with nandrolone (15 mg/kg) for 30 days showed reduced locomotor activity in an open field and EPM (El-Shamarka et al., 2020). In adults, nandrolone is reported to decrease activity in the running wheel (Keleta et al., 2007), in an open field (Rosic et al., 2014; Selakovic et al., 2017) and in the EPM (Minkin et al., 1993). Nonetheless there are some studies that fail to find an effect (Salvador et al., 1999), and others that report a decrease in cocaine-induced locomotor activity (Kurling-Kailanto et al, 2010) after nandrolone withdrawal.

The mechanism by which nandrolone decreases locomotor activity is not clear. Purves-Tyson et al (2012) have reported that testosterone administration to prepubertal rats modulates the expression of enzymes that participate in dopaminergic metabolism in brain regions, such as the substantia nigra, that regulate locomotion and affect. Others report that the addictive properties of AAS may be exerted through the endogenous opioid system that in turn stimulate dopaminergic centers in the brain (Bontempi and Bonci, 2020).

### 5.4 AAS and cocaine

Several studies indicate that AAS users are likely to develop a dependence on other drugs of abuse (Mȩdraś et al., 2018; Pope et al., 2014a). These results must be interpreted with caution since, in some cases, AAS users abused other drugs prior to using AAS. In addition, studies in adult rodents investigating the role of androgens in modulating the response to drugs of abuse have provided conflicting reports.

Prior studies from our laboratory and of others indicate that removal of the primary source of androgens by gonadectomy increases the response to psychostimulants (Beatty et al., 1982; Dluzen et al., 1986; Forgie and Wart, 1994; Menéndez-Delmestre and Segarra, 2011; Purves-Tyson et al., 2015), although not all studies have obtained similar results (Caine et al., 2004; Chen et al., 2003; Chin et al., 2002; Haney et al., 1994; Harrod et al., 2005; Jackson et al., 2006). Furthermore, testosterone administration to adult males is reported to decrease the acute locomotor response to cocaine and amphetamine in gonadectomized (Menéndez-Delmestre and Segarra, 2011; Purves-Tyson et al., 2015) and gonadally intact drug-naive males (Beatty et al., 1982; Long et al., 1994). Indeed, our laboratory has found that testosterone is necessary for adult male rats to develop and express sensitization to cocaine (Menéndez-Delmestre and Segarra, 2011).

Several studies indicate that prepubertal male rats do not become sensitized to cocaine at this early age (Kabbaj et al., 2002; Lepsch et al., 2005; Trzcińska et al., 2002). Indeed, our control rats do not show sensitization until after a withdrawal period and re-exposure to cocaine. Oil-cocaine rats displayed sensitization when they were 52 days of age. At this time, they had received 5 daily cocaine injections, a 7-day withdrawal period and a cocaine challenge (day 13 of test). Sensitization in these oil-treated animals became more robust after a second withdrawal period and re-exposure to cocaine (day 23), when they were 62 days of age (Fig 4C). In contrast, we found that exposure to nandrolone during days 28-37 accelerated the development of sensitization, which was evident after 5 cocaine injections, when they are 44 days of age (Fig 4D).

We also observed that the hyperactivity induced by cocaine is maintained for a longer time period in nandrolone-treated males during the first cocaine challenge (day 13). It is unclear if this effect can be attributed to changes in the metabolism of cocaine. Cocaine and nandrolone are metabolized mainly in the liver, cocaine by esterases and cytochrome P450 enzymes, whereas metabolism of nandrolone occurs mainly by 5α-reductase and 3α- and 3β-hydroxysteroid dehydrogenase enzymes. Thus, it is not surprising that pharmacokinetic interactions between the compounds have been observed. Synergistics effects on the cardiovascular system (Tseng et al, 1994) and seizures (Long et al (2000), as well as alterations in brain dopaminergic and serotonergic outflow (Kurling-Kailanto et al, 2010), have been reported. Unfortunately, although areas of the mesocorticolimbic pathway are rich in androgen receptors (Tobiansky et al., 2018), very few studies have investigated if androgens alter the metabolism of cocaine (Bowman et al., 1999; Lukas et al., 1996). Another possibility is that exposure to androgens during the prepubertal period accelerates the maturation of some component of the mesocorticolimbic circuitry essential for the display of behavioral sensitization. For example, the cytochrome P450 system is involved in the metabolism of androgens and cocaine (McDonnell and Dang, 2013); thus, both drugs may interact pharmacokinetically and affect the individual response of each drug. A study by Yamamoto et al (2007) indicates that treatment with the antiandrogen flutamide decreases plasma levels of cocaine and its main metabolites, suggesting that testosterone may potentiate the effects of cocaine, as we have previously reported (Menéndez-Delmestre and Segarra, 2011).

Our data also indicates that prior exposure to AAS increases the behavioral response to cocaine, a process known as cross sensitization. Cross sensitization occurs with previous exposure between the same drug and also between different drugs of abuse (Smith et al., 2013). The mechanisms that mediate cross sensitization are not fully understood. Evidence suggests that dopamine is the substrate involved in this process (Kalivas and Stewart, 1991). Studies found that androgens like testosterone induce cross sensitization to cocaine in prepubertal, but not in adult male rats (Engi et al., 2015).

One scenario may be that AAS promotes cross sensitization by activating gene expression of corticotropin releasing hormone (CRH). The gene that codes for CRH contains androgen and estrogen response elements that modulate expression of CRH (Bao et al., 2006; Bao and Swaab, 2007). Indeed, we have recently reported that altered corticotropin-releasing hormone receptor 1 (CRF-R1) sensitivity may lead to the observed DA hyperresponsiveness observed in socially isolated adolescent rats (Novoa et al., 2021).

### 5.5 CPP

All rats tested between days 40 to 53 days developed CPP to cocaine. In addition, oil-cocaine rats spent more time in the chamber associated with cocaine during the post-conditioning day compared to nandrolone-treated rats. The increase in time spent in the chamber associated with cocaine was 124% in oil-cocaine rats compared to 57% in those pretreated with nandrolone. Thus, although both groups of rats showed CPP to cocaine, CPP was lower in rats that previously received nandrolone.

Our studies agree with that of others that report a decrease in CPP to other drugs of abuse such as cannabinoids (Celerier et al., 2006) and opioids (Célérier et al. 2003) following AAS administration. In addition, exposure to nandrolone (15 mg/kg) during PN 40-53 decreased sucrose consumption (Rainer et al, 2014) 15 days after withdrawal. Nonetheless, there are studies that indicate that withdrawal from nandrolone increases cannabinoids (Struik et al., 2016) and alcohol (Johansson et al. 2000) intake. Thus, there is no consensus indicating if prior exposure to nandrolone results in an enhancement or an aversion to other drugs of abuse or rewarding stimuli such as sucrose. However, there is evidence that the addictive properties of AAS may be exerted through the endogenous opioid system that in turn stimulate dopaminergic centers in the brain (Bontempi and Bonci, 2020).

The CPP paradigm involves several cognition components such as acquisition, retrieval, and extinction of spatial and contextual memories (Anagnostaras et al., 1999; Gould and Leach, 2014; Riedel et al., 1999). Several lines of evidence link the use of AAS with cognitive dysfunction (Wallin and Wood, 2015) and altered decision-making when studied in paradigms such as the operant discounting task (Cooper et al., 2014; Wallin-Miller et al., 2018; Wallin et al., 2015; Wood et al., 2013).

We cannot discard the possibility that during the CPP test, the reduced time spent in the chamber associated with cocaine is due to a decrease in motivation. However, the increased sensitized locomotor response to cocaine displayed at day 5 by animals that received nandrolone elicit uncertainty on this possibility. Sensitization has been defined as the successive augmentation of locomotor hyperactivity elicited by repeated administration of psychostimulants. It involves neuroadaptations in the mesocorticolimbic system that contribute to changes in the motivational circuitry underlying craving and relapse (Chefer et al., 2005; Kumar et al., 2005; Thomas et al., 2008). Since many investigators relate sensitization with increased motivation, and nandrolone-treated animals displayed increased sensitization, we are inclined to favor alterations in cognitive, and not rewarding, aspects of CPP. In addition, the rewarding properties of AAS are evident in previous studies that find that rodents will self-administer AAS orally and intracranially (Johnson and Wood, 2001; Wood, 2004), an effect that disappears with administration of an androgen receptor antagonist (Peters and Wood, 2005).

### 5.6 AAS and D2DR

At the conclusion of the sensitization and CPP experiments, we investigated if D2DR receptors in the NAc and mPFC were affected by prior treatment with nandrolone, cocaine or both. Separate groups of animals were used for the above-mentioned experiments. Rats used for CPP received 5 cocaine injections (15 mg/kg) one every other day for 10 days and killed 24 hrs after the last cocaine injection. Rats used for the sensitization experiments received 7 cocaine injections (15 mg/kg) during a 23 day period: 5 daily injections, a 7 day withdrawal period, 1 challenge injection, a 9 day withdrawal period, a second challenge injection and euthanized 24 hours after the last cocaine injection. Thus, the sensitization group received two additional cocaine injections and underwent two drug-free periods of 7 and 9 days prior to euthanasia.

### 5.7 D2DR in the NAc of rats used for the sensitization experiments

Rats treated with nandrolone (days 28-37) had a higher concentration of D2DR in the NAc at the day of euthanasia (days 54 and 63) compared to oil-saline rats. In contrast, all groups that received cocaine had lower levels of D2DR compared to oil saline rats. Interestingly, groups that were injected with nandrolone (days 28-37) and used for the sensitization experiments (euthanized at day 63) had the lowest levels of D2DR in the NAc of all groups.

Behavioral sensitization has two phases, initiation, and expression. The initiation comprises rapid neural effects that induce behavioral sensitization, the expression involves the long-term consequences (Kalivas and Stewart, 1991). Initiation of behavioral sensitization does not require activation of dopamine receptors (White et al., 1998), however its expression does (Steketee and Kalivas, 2011; Thomas et al., 2001; Vanderschuren and Kalivas, 2000). Most studies agree that the NAc, although not essential for the development of locomotor sensitization to cocaine, is necessary for its expression (Di Chiara, 1995). Unfortunately, the role of each dopaminergic receptor subtype in the process of sensitization is still not entirely clear.

Some studies report that D2DR in the NAc do not affect cocaine-induced locomotor activity nor behavioral sensitization (Beyer and Steketee, 2002; Mattingly et al., 1994; Vanderschuren and Kalivas, 2000). However, other studies attest to D2DRs modulation of cocaine-induced sensitization (Di Chiara, 1995). For example, blocking D2DR receptors in the NAc significantly decreased cocaine-induced locomotor activity (Baker et al., 1996; Manvich et al., 2019; Neisewander et al., 1998) and abolished cocaine-induced sensitization (Manvich et al, 2019). Furthermore, deletion of D2DR in medium spiny neurons (MSN) of the NAc in mice results in decreased cocaine-induced locomotor activity (Smith et al., 2013). In contrast, experiments using the conditional mutant mice “autodrd2KO”, which is characterized by a lack of D2DR auto receptors (those in dopamine neurons and terminals), indicate that sensitivity to cocaine is enhanced in these animals (Bello et al., 2011). Mice lacking D2DR auto receptors also show enhanced CPP as well as hyperlocomotion (without altering dopamine transporters function) (Bello et al, 2011).

These data argue that the process by which D2DRs in the NAc modulate the response to cocaine can vary depending if these receptors are located on dopaminergic terminals or GABAergic MSN in the NAc and may partly explain several conflicting reports. It is possible that the observed decrease in D2DR in the NAc in this study occurred mainly in dopaminergic terminals. Repeated exposure to cocaine during sensitization would transiently result in increased dopamine (DA) that in turn would induce D2DR downregulation. D2DRs have a greater affinity for DA than D1DR (Dreyer et al., 2010; Rice and Cragg, 2008). This would result in less dopaminergic autoinhibition and greater, or extended, DA release, resulting in an exacerbated locomotor response to cocaine, as we observed in the current study. It is also possible that DA, by modulating AMPA trafficking, contributes to the enhancement of sensitization (Boudreau and Wolf, 2005). However, more studies are needed to determine the cell type or terminal where the decrease in D2DR occurred.

In addition, nandrolone treatment increased D2DR in the NAc. This group of animals displayed the highest locomotor activity in response to cocaine, and developed sensitization earlier than oil-treated rats, (after 5 cocaine injections). These data are in accordance with previous studies that show that D2DR availability predicts future drug-seeking behavior (Haney et al., 1994)

### 5.8 Androgens modulate D2DR

Androgens can modulate dopamine receptors in the mesocorticolimbic circuitry (Aubele and Kritzer, 2012; Bertozzi et al., 2018; Purves-Tyson et al., 2012; Tobiansky et al., 2018). Kindlundh et al (2001) found that administering nandrolone for two weeks at low doses (1 and 5 mg/kg) increased D2DR in the NAc core and shell of male rats but found that a higher dose (15 mg/kg) had no effect. The authors of this last study did not state the age, nor weight, of the rats. In our study, rats were 28 days of age when they received the first nandrolone injection, and the dose used was 20 mg/kg. Thus, we cannot determine if the difference between these two studies is dose or age-related.

Nandrolone can also alter DA metabolism (Zotti et al., 2014). A decrease in the activity of the dopamine-metabolizing enzymes monoamine oxidase A and B has been reported after nandrolone treatment (Birgner et al., 2008, 2007). Furthermore, levels of tyrosine hydroxylase in the substantia nigra increase prior to puberty, coinciding with the increase in testosterone (Purves -Tyson et al., 2012).

The Akil group, among others, have selectively bred rats according to their initial response to a novel environment and classified them as high responders (HR) and low responders (LR) (Aydin et al., 2021; Birt et al., 2021; Clinton et al., 2012; Turner et al., 2009). These two lines also show differences in behavioral traits relevant to addiction, with HR displaying a greater amount of drug taken, persistence of drug-seeking and drug-induced locomotor activity (Flagel et al., 2016). They also differ in the neural substrates that regulate addictive behaviors. Like the nandrolone-treated males in this study, HRs have higher D2DR in the NAc core compared to LR (Clinton et al., 2012). HRs also have greater fibroblast growth factor 2 (FGF2) and lower D1DR levels in the NAc. Interestingly, previous studies show that testosterone increases plasma levels of FGF2 and of Insulin Growth Factor (Ghanim et al., 2019) and mRNA expression of FGF2 *in vitro* (Saito et al., 1991). Dysregulation of neurotrophic factors like FGF2 is involved in increased vulnerability to drugs (McGinty et al., 2010; Thomas et al., 2008). Indeed, drugs such as amphetamines and cocaine modulate the expression of FGF2 (Mueller et al., 2006; Turner et al., 2009). This may explain the synergistic effect of nandrolone and cocaine in decreasing D2DR in the NAc.

### 5.9 Prefrontal cortex

The mPFC plays an important role in addictive behaviors such as decision making, memory retrieval and cocaine seeking behaviors and is necessary for the induction of sensitization to cocaine (Cador et al., 1999; Li et al., 1999; Dalley et al., 2004; Moorman et al., 2015). It contains a distinct population of glutamatergic pyramidal neurons that project to the striatum and other subcortical regions that express D1DR and D2DR (Gaspar et al., 1995). Dopaminergic modulation of glutamatergic function contributes to reward, salience, attention, and working memory (Brozoski et al., 1979; Chudasama and Robbins, 2004; Lohani et al., 2019; Yokel and Wise, 1975). Recent evidence indicates that DA modulates ensemble activity, facilitating and strengthening information processing in the PFC (Lohani et al., 2019).

Maturation of PFC circuitry continues after puberty, dopaminergic innervation from the VTA to the PFC increases gradually until approximately PN day 60 (Salas et al., 2016). At approximately this age, males have acquired adult testosterone plasma levels and the ability to display male sexual behavior (Segarra and Strand, 1989). An increase in dendritic spine density following androgen or estrogen administration suggest that gonadal steroids play a role in PFC function (Hajszan et al., 2007), Interestingly, on day 40, an increase in D2DR receptors in the PFC is observed, coinciding with the activation of the HPG axis in male rats (Andersen et al., 2000). It is possible that in our current study, that administration of nandrolone from day 28 to 37 accelerated maturation of the motivational circuitry in the PFC, which in turn could be responsible for the expression of sensitization at a younger age.

### Summary

In summary, this study provides evidence that exposure to the AAS nandrolone prepubertally increases risk-taking behaviors and decreases anxiety and locomotor activity. Nandrolone also exacerbated the locomotor response to cocaine and accelerated the development of cocaine sensitization, occurring at an earlier age and with less cocaine injections. It highlights the role of androgens in accelerating the development of brain structures that participate in the psychomotor response to drugs of abuse, as illustrated by the display of cocaine-induced behavioral sensitization in nandrolone-treated prepubertal males. This study also confirms that the effect of AAS administration prepubertally on the remodeling of neural circuitry may occur quite rapidly after exposure. Changes in accumbal dopaminergic receptors hint at androgenic modulation of mesocorticolimbic substrates during early adolescent development. The high prevalence of AAS abuse worldwide and the long-lasting and detrimental effects on essential physiological processes and neural circuitries that regulate addictive and social behaviors warrants further studies.

**Table 4.**
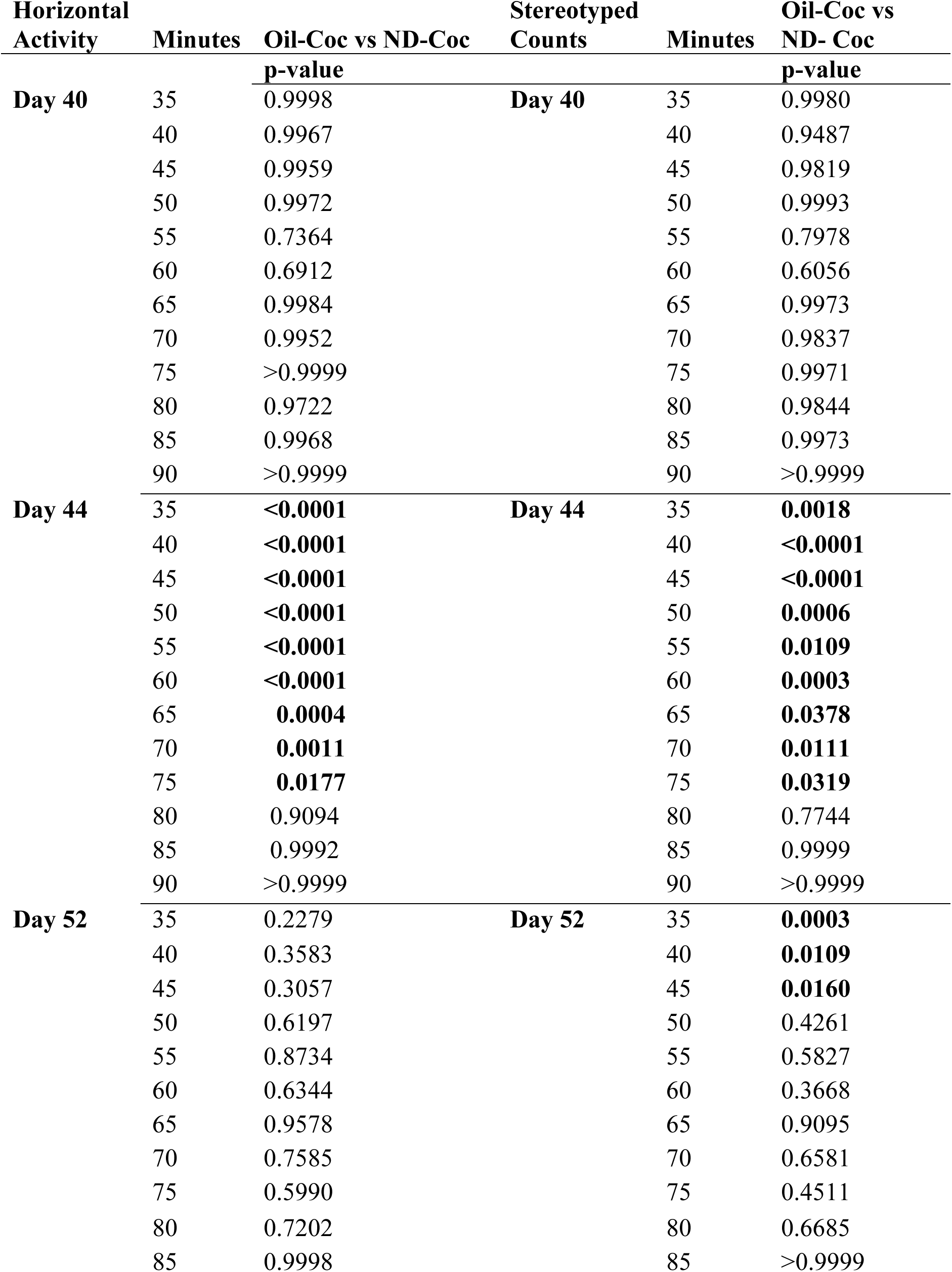
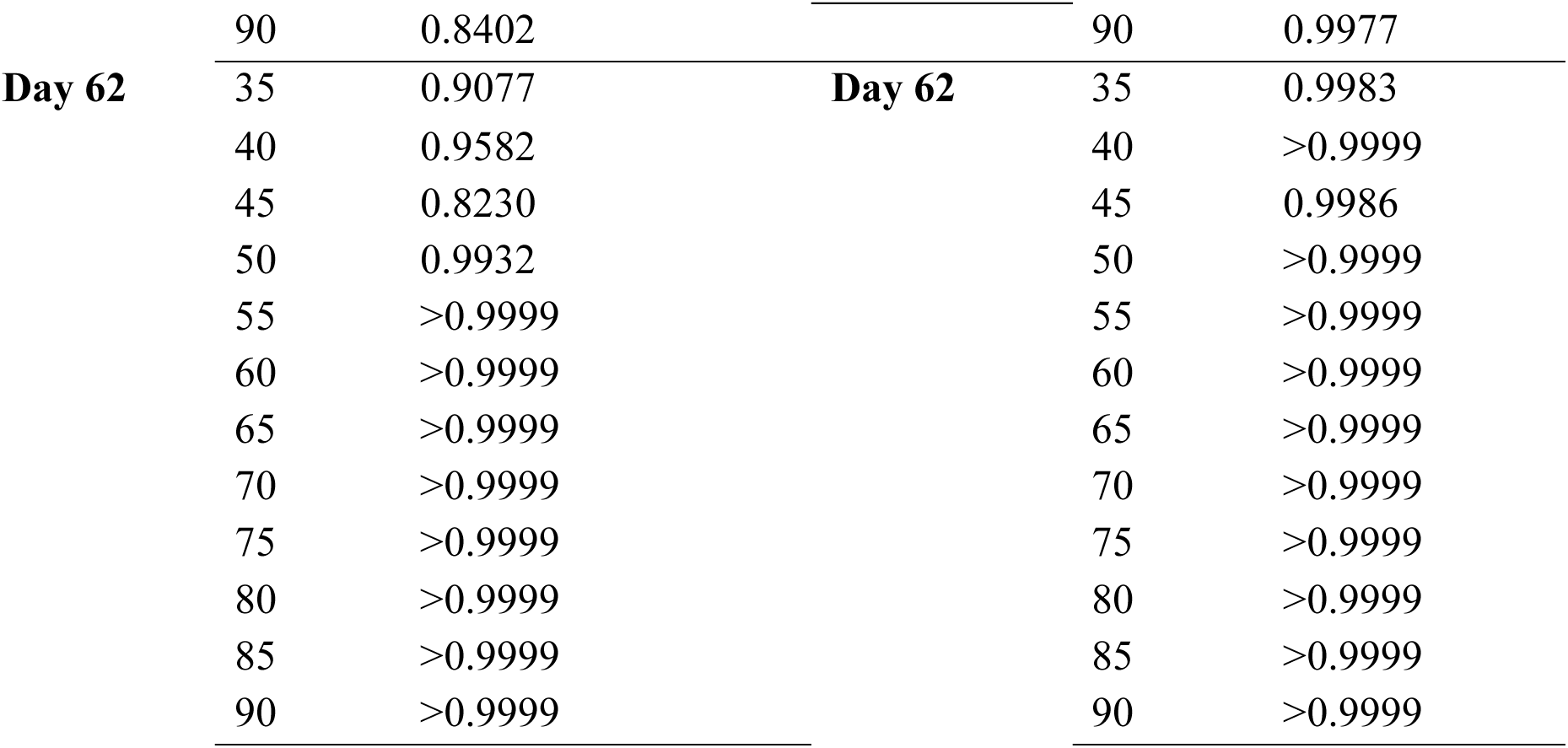
Two-Way ANOVA Post-hoc Analysis Oil-Coc vs ND-Coc.

## Acknowledgements

The authors would like to thank Lidalee Silva, Erick Quintana and Bethzaida Suarez for their technical assistance.

## REFERENCES

Aikey, J.L., Nyby, J.G., Anmuth, D.M., James, P.J., 2002. Testosterone rapidly reduces anxiety in male house mice (Mus musculus). Horm. Behav. 42, 448–460. 10.1006/hbeh.2002.1838

Anagnostaras, S.G., Maren, S., Fanselow, M.S., 1999. Temporally graded retrograde amnesia of contextual fear after hippocampal damage in rats: Within-subjects examination. J. Neurosci, 19(3), 1106–1114. 10.1523/jneurosci.19-03-01106.1999

Andersen, S.L., Thompson, A.T., Rutstein, M., Hostetter, J.C., Teicher, M.H., 2000.Dopamine receptor pruning in prefrontal cortex during the periadolescent period in rats. Synapse, 37(2), 167–169. 10.1002/1098-2396(200008)37:2<167::AID-SYN11>3.0.CO;2-B

Aydin, C., Frohmader, K., Emery, M., Blandino, P., Akil, H., 2021. Chronic stress in adolescence differentially affects cocaine vulnerability in adulthood in a selectively bred rat model of individual differences: role of accumbal dopamine signaling. Stress, 24(3), 251–260. 10.1080/10253890.2020.1790520

Aubele, T., Kritzer, M.F., 2012. Androgen influence on prefrontal dopamine systems in adult male rats: Localization of cognate intracellular receptors in medial prefrontal projections to the ventral tegmental area and effects of gonadectomy and hormone replacement on glutamate stimulated. Cereb. Cortex 22, 1799–1812. 10.1093/cercor/bhr258

Bahrke, M.S., Yesalis, C.E., Wright, J.E., 1996. Psychological and behavioural effects of endogenous testosterone and anabolic-androgenic steroids. An update. Sport, 22(6), 367–390. Med. 10.2165/00007256-199622060-00005

Basaria, S., Dobs, A.S., 2001. Hypogonadism and androgen replacement therapy in elderly men. Am. J. Med, , 110(7), 563–572. 10.1016/S0002-9343(01)00663-5

Beatty, W.W., Dodge, A.M., Traylor, K.L., 1982. Stereotyped behavior elicited by amphetamine in the rat: Influences of the testes. Pharmacol. Biochem. Behav, 16(4), 565–568. 10.1016/0091-3057(82)90416-6

Bello, E.P., Mateo, Y., Gelman, D.M., Noaín, D., Shin, J.H., Low, M.J., Alvarez, V.A., Lovinger, D.M., Rubinstein, M., 2011. Cocaine supersensitivity and enhanced motivation for reward in mice lacking dopamine D2 autoreceptors. Nat. Neurosci, 14(8), 1033–1038. 10.1038/nn.2862

Bergink, E.W., Janssen, P.S.L., Turpun, E.W., Van Der Vies, J., 1985. Comparison of the receptor binding properties of nandrolone and testosterone under in vitro and in vivo conditions. J. Steroid Biochem, 22(6), 831–836. 10.1016/0022-4731(85)90293-6

Bertozzi, G., Sessa, F., Albano, G.D., Sani, G., Maglietta, F., Roshan, M.H.K., Volti, G.L., Bernardini, R., Avola, R., Pomara, C., Salerno, M., 2018. The Role of Anabolic Androgenic Steroids in Disruption of the Physiological Function in Discrete Areas of the Central Nervous System. Mol. Neurobiol, 55(7), 5548–5556. 10.1007/s12035-017-0774-1

Beyer, C.E., Steketee, J.D., 2002. Cocaine sensitization: Modulation by dopamine D2 receptors. Cereb. Cortex, 12(5), 526–535. 10.1093/cercor/12.5.526

Biedermann, S. V., Biedermann, D.G., Wenzlaff, F., Kurjak, T., Nouri, S., Auer, M.K., Wiedemann, K., Briken, P., Haaker, J., Lonsdorf, T.B., Fuss, J., 2017. An elevated plus-maze in mixed reality for studying human anxiety-related behavior. BMC Biol, 15(1). 10.1186/s12915-017-0463-6

Birgner, C., Kindlundh-Högberg, A.M.S., Alsiö, J., Lindblom, J., Schiöth, H.B., Bergström, L., 2008. The anabolic androgenic steroid nandrolone decanoate affects mRNA expression of dopaminergic but not serotonergic receptors. Brain Res, 1240, 221–228. 10.1016/j.brainres.2008.09.003

Birgner, C., Kindlundh-Högberg, A.M.S., Nyberg, F., Bergström, L., 2007. Altered extracellular levels of DOPAC and HVA in the rat nucleus accumbens shell in response to sub-chronic nandrolone administration and a subsequent amphetamine challenge. Neurosci. Lett, 412(2), 168–172. 10.1016/j.neulet.2006.11.001

Birt, I.A., Hagenauer, M.H., Clinton, S.M., Aydin, C., Blandino, P., Stead, J.D.H., Hilde, K.L., Meng, F., Thompson, R.C., Khalil, H., Stefanov, A., Maras, P., Zhou, Z., Hebda-Bauer, E.K., Goldman, D., Watson, S.J., Akil, H., 2021. Genetic Liability for Internalizing Versus Externalizing Behavior Manifests in the Developing and Adult Hippocampus: Insight From a Meta-analysis of Transcriptional Profiling Studies in a Selectively Bred Rat Model. Biol. Psychiatry, 89(4), 339–355. 10.1016/j.biopsych.2020.05.024

Bitran, D., Kellogg, C.K., Hilvers, R.J., 1993. Treatment with an Anabolic-Androgenic Steroid Affects Anxiety-Related Behavior and Alters the Sensitivity of Cortical GABAA Receptors in the Rat. Horm. Behav, 27(4), 568–583. 10.1006/hbeh.1993.1041

Boudreau, A.C., Wolf, M.E., 2005. Behavioral sensitization to cocaine is associated with increased AMPA receptor surface expression in the nucleus accumbens. J. Neurosci, 25(40), 9144–9151. 10.1523/JNEUROSCI.2252-05.2005

Bowman, B.P., Vaughan, S.R., Walker, Q.D., Davis, S.L., Little, P.J., Scheffler, N.M., Thomas, B.F., Kuhn, C.M., 1999. Effects of sex and gonadectomy on cocaine metabolism in the rat. J. Pharmacol. Exp. Ther, 290(3), 1316–1323.

Brower, K.J., Blow, F.C., Beresford, T.P., Fuelling, C., 1989. Anabolic-androgenic steroid dependence. J. Clin. Psychiatry, 50(1), 31–33.

Brower, K.J., Eliopulos, G.A., Blow, F.C., Catlin, D.H., Beresford, T.P., 1990. Evidence for physical and psychological dependence on anabolic androgenic steroids in eight weight lifters. Am. J. Psychiatry, 147(4), 510–512. 10.1176/ajp.147.4.510

Brozoski, T.J., Brown, R.M., Rosvold, H.E., Goldman, P.S., 1979. Cognitive deficit caused by regional depletion of dopamine in prefrontal cortex of rhesus monkey. Science, 205(4409), 929–932. 10.1126/science.112679

Bruchovsky, N., Wilson, J.D., 1999. Discovery of the role of dihydrotestosterone in androgen action. Steroids, 64(11), 753–759. 10.1016/S0039-128X(99)00054-9

Cador, M., Bjijou, Y., Cailhol, S., Stinus, L., 1999. D-Amphetamine-induced behavioral sensitization: Implication of a glutamatergic medial prefrontal cortex-ventral tegmental area innervation. Neuroscience, 94(3), 705–721.

Caine, S.B., Bowen, C.A., Yu, G., Zuzga, D., Negus, S.S., Mello, N.K., 2004. Effect of gonadectomy and gonadal hormone replacement on cocaine self-administration in female and male rats. Neuropsychopharmacology, 29(5), 929–942. 10.1038/sj.npp.1300387

Célérier E, Yazdi MT, Castañé, A, Ghozland S, Nyberg F and R Maldonado 2003. Effects of nandrolone on acute morphine responses, tolerance and dependence in mice. Eur J Pharmacol 465 (1-2):69–81. doi: 10.1016/s0014-2999(03)01462-6.

Célérier E, Ahdepil T, Wikander H, Berrendero F, Nyberg F and R Maldonado 2006. Influence of the anabolic-androgenic steroid nandrolone on cannabinoid dependence. Neuropharmacol 50 (7):788–806. doi: 10.1016/j.neuropharm.2005.11.017. Epub 2006 Jan 27.

Chen, R., Osterhaus, G., McKerchar, T., Fowler, S.C., 2003. The role of exogenous testosterone in cocaine-induced behavioral sensitization and plasmalemmal or vesicular dopamine uptake in castrated rats. Neurosci. Lett, 351(3), 161–164. 10.1016/j.neulet.2003.07.018

Chin, J., Sternin, O., Wu, H., Burrell, S., Lu, D., Jenab, S., Perrotti, L.I., Quiñones-Jenab, V., 2002. Endogenous gonadal hormones modulate behavioral and neurochemical responses to acute and chronic cocaine administration. Brain Res, 945(1), 123–130. 10.1016/S0006-8993(02)02807-X

Chudasama, Y., Robbins, T.W., 2004. Dopaminergic modulation of visual attention and working memory in the rodent prefrontal cortex. Neuropsychopharmacology, 29(9), 1628–1636. 10.1038/sj.npp.1300490

Clinton, S.M., Turner, C.A., Flagel, S.B., Simpson, D.N., Watson, S.J., Akil, H., 2012. Neonatal fibroblast growth factor treatment enhances cocaine sensitization. Pharmacol. Biochem. Behav, 103(1), 6–17. 10.1016/j.pbb.2012.07.006

Cooper, S.E., Goings, S.P., Kim, J.Y., Wood, R.I., 2014. Testosterone enhances risk tolerance without altering motor impulsivity in male rats. Psychoneuroendocrinology, 40, 201–212.

Dalley, J.W., Theobald, D.E., Bouger, P., Chudasama, Y., Cardinal, R.N., Robbins, T.W., 2004. Cortical cholinergic function and deficits in visual attentional performance in rats following 192 IgG-saporin-induced lesions of the medial prefrontal cortex. Cereb. Cortex, 14(8), 922–932. 10.1093/cercor/bhh052

Denham, B.E., 2006. Effects of mass communication on attitudes toward anabolic steroids: An analysis of high school seniors. J. Drug Issues, 36(4), 809–829. 10.1177/002204260603600403

Di Chiara, G., 1995. The role of dopamine in drug abuse viewed from the perspective of its role in motivation. Drug Alcohol Depend, 38(2), 95–137. 10.1016/0376-8716(95)01118-I

Dluzen, D.E., Green, M.A., Ramirez, V.D., 1986. The effect of hormonal condition on dose-dependent amphetamine-stimulated behaviors in the male rat. Horm. Behav, 20(1), 1–6. 10.1016/0018-506X(86)90024-3

Dokras, A., Clifton, S., Futterweit, W., Wild, R., 2012. Increased prevalence of anxiety symptoms in women with polycystic ovary syndrome: Systematic review and meta-analysis. Fertil. Steril, 97(1), 225–230.e2. 10.1016/j.fertnstert.2011.10.022

Dreyer, J.K., Herrik, K.F., Berg, R.W., Hounsgaard, J.D., 2010. Influence of phasic and tonic dopamine release on receptor activation. J. Neurosci, 30(42), 14273–14283. 10.1523/JNEUROSCI.1894-10.2010

DuRant, R.H., Escobedo, L.G., Heath, G.W., 1995. Anabolic-steroid use, strength training, and multiple drug use among adolescents in the United States. , 94(3), 705–721. 10.1016/s0306-4522(99)00361-9

Edinger, K.L., Frye, C.A., 2006. Intrahippocampal administration of an androgen receptor antagonist, flutamide, can increase anxiety-like behavior in intact and DHT-replaced male rats. Horm. Behav, 50(2), 216–222. 10.1016/j.yhbeh.2006.03.003

El-Shamarka, M.E.S., Sayed, R.H., Assaf, N., Zeidan, H.M., Hashish, A.F., 2020. Combined neurotoxic effects of cannabis and nandrolone decanoate in adolescent male rats. Neurotoxicology, 76, 114–125. 10.1016/j.neuro.2019.11.001

Engi, S.A., Cruz, F.C., Crestani, C.C., Planeta, C.S., 2015. Cross-sensitization between testosterone and cocaine in adolescent and adult rats. Int. J. Dev. Neurosci, 46(1), 33–37. 10.1016/j.ijdevneu.2015.07.001

Flagel, S.B., Chaudhury, S., Waselus, M., Kelly, R., Sewani, S., Clinton, S.M., Thompsona, R.C., Watson, S.J., Akila, H., 2016. Genetic background and epigenetic modifications in the core of the nucleus accumbens predict addiction-like behavior in a rat model. Proc. Natl. Acad. Sci, 113(20), E2861–E2870. 10.1073/pnas.1520491113

Forgie, M.L., Stewart, J., 1994. Sex differences in the locomotor-activating effects of amphetamine: Role of circulating testosterone in adulthood. Physiol. Behav, 55(4), 639–644. 10.1016/0031-9384(94)90038-8

Gaspar, P., Bloch, B., Le Moine, C., 1995. D1 and D2 Receptor Gene Expression in the Rat Frontal Cortex: Cellular Localization in Different Classes of Efferent Neurons. Eur. J. Neurosci, 7(5), 1050–1063. 10.1111/j.1460-9568.1995.tb01092.x

Ghanim, H., Dhindsa, S., Batra, M., Green, K., Abuaysheh, S., Kuhadiya, N.D., Makdissi, A., Chaudhuri, A., Dandona, P., 2019. Effect of testosterone on FGF2, MRF4, and myostatin in hypogonadotropic hypogonadism: Relevance to muscle growth. J. Clin. Endocrinol. Metab, 104(6), 2094–2102. 10.1210/jc.2018-01832

Ghods-Sharifi, S., Floresco, S.B., 2010. Differential effects on effort discounting induced by inactivations of the nucleus accumbens core or shell. Behav. Neurosci, 124(2), 179–191. 10.1037/a0018932

Gould, T.J., Leach, P.T., 2014. Cellular, molecular, and genetic substrates underlying the impact of nicotine on learning. Neurobiol. Learn. Mem, 107, 108–132. 10.1016/j.nlm.2013.08.004

Grönbladh, A., Nylander, E., Hallberg, M., 2016. The neurobiology and addiction potential of anabolic androgenic steroids and the effects of growth hormone. Brain Res. Bull. 126, 127–137. 10.1016/j.brainresbull.2016.05.003

Hajszan, T., MacLusky, N.J., Johansen, J.A., Jordan, C.L., Leranth, C., 2007. Effects of androgens and estradiol on spine synapse formation in the prefrontal cortex of normal and testicular feminization mutant male rats. Endocrinology, 148(5), 1963– 1967. 10.1210/en.2006-1626

Haney, M., Castanon, N., Cador, M., Le Moal, M., Mormède, P., 1994. Cocaine sensitivity in roman high and low avoidance rats is modulated by sex and gonadal hormone status. Brain Res, 645(1–2), 179–185. 10.1016/0006-8993(94)91651-9

Harrod, S.B., Mactutus, C.F., Browning, C.E., Welch, M., Booze, R.M., 2005. Home cage observations following acute and repeated IV cocaine in intact and gonadectomized rats. Neurotoxicol. Teratol, 27(6), 891–896. 10.1016/j.ntt.2005.07.004

Hosking, J.G., Floresco, S.B., Winstanley, C.A., 2015. Dopamine antagonism decreases willingness to expend physical, but not cognitive, effort: A comparison of two rodent cost/benefit decision-making tasks. Neuropsychopharmacology, 40(4), 1005–1015. 10.1038/npp.2014.285

Hu, M., Richard, J.E., Maliqueo, M., Kokosar, M., Fornes, R., Benrick, A., Jansson, T., Ohlsson, C., Wu, X., Skibicka, K.P., Stener-Victorin, E., 2015. Maternal testosterone exposure increases anxiety-like behavior and impacts the limbic system in the offspring. Proc. Natl. Acad. Sci, 112(46), 14348–14353. 10.1073/pnas.1507514112

Hyman, S.E., Malenka, R.C., Nestler, E.J., 2006. Neural mechanisms of addiction: The role of reward-related learning and memory. Annu. Rev. Neurosci, 29(1), 565–598. 10.1146/annurev.neuro.29.051605.113009

Jackson, L.R., Robinson, T.E., Becker, J.B., 2006. Sex differences and hormonal influences on acquisition of cocaine self-administration in rats. Neuropsychopharmacology, 31(1), 129–138. 10.1038/sj.npp.1300778

Johnson, L.R., Wood, R.I., 2001. Oral testosterone self-administration in male hamsters. Neuroendocrinology, 73(4), 285–292. 10.1159/000054645

Kabbaj, M., Isgor, C., Watson, S.J., Akil, H., 2002. Stress during adolescence alters behavioral sensitization to amphetamine. Neuroscience, 113(2), 395–400. 10.1016/S0306-4522(02)00188-4

Kalivas, P.W., Stewart, J., 1991. Dopamine transmission in the initiation and expression of drug- and stress-induced sensitization of motor activity. Brain Res. Rev, 16(3), 223–244. 10.1016/0165-0173(91)90007-U

Kanayama, G., Hudson, J.I., Pope, H.G., 2008. Long-term psychiatric and medical consequences of anabolic-androgenic steroid abuse: A looming public health concern? Drug Alcohol Depend, 98(1–2), 1–12. 10.1016/j.drugalcdep.2008.05.004

Kaufman, M.J., Janes, A.C., Hudson, J.I., Brennan, B.P., Kanayama, G., Kerrigan, A.R., Jensen, J.E., Pope, H.G., 2015. Brain and cognition abnormalities in long-term anabolic-androgenic steroid users. Drug Alcohol Depend, 152, 47–56. 10.1016/j.drugalcdep.2015.04.023

Keleta, Y.B., Lumia, A.R., Anderson, G.M., McGinnis, M.Y., 2007. Behavioral effects of pubertal anabolic androgenic steroid exposure in male rats with low serotonin. Brain Res. 1132, 129–138. 10.1016/j.brainres.2006.10.097

Kicman, A.T., 2008. Pharmacology of anabolic steroids. Br. J. Pharmacol. 154, 502–521. 10.1038/bjp.2008.165

Kindlundh, A.M.S., Isacson, D.G.L., Berglund, L., Nyberg, F., 1999. Factors associated with adolescent use of doping agents: Anabolic-androgenic steroids. Addiction, 94(4), 543–553. 10.1046/j.1360-0443.1999.9445439.x

Kindlundh, Anna M.S., Lindblom, J., Bergström, L., Wikberg, J.E.S., Nyberg, F., 2001. The anabolic-androgenic steroid nandrolone decanoate affects the density of dopamine receptors in the male rat brain. Eur. J. Neurosci, 13(2), 291–296. 10.1046/j.0953-816X.2000.01402.x

Kochakian, C.D., Welder, A.A., 1993. Anabolic-androgenic steroids: In cell culture. Vitr. Cell. Dev. Biol. - Anim, 29(6), 433–438. 10.1007/BF02639373

Kouvelas, D., Pourzitaki, C., Papazisis, G., Dagklis, T., Dimou, K., Kraus, M.M., 2008. Nandrolone abuse decreases anxiety and impairs memory in rats via central androgenic receptors. Int. J. Neuropsychopharmacol, 11(07). 10.1017/S1461145708008754

Kurling-Kailanto S, Kankaanpää, A, Seppälä T 2010. Subchronic nandrolone administration reduces cocaine-induced dopamine and 5-hydroxytryptamine outflow in the rat nucleus accumbens. Psychopharmacology, v. 209, p. 271–281. doi: 10.1007/s00213-010-1796-9. Epub 2010 Feb 26.

Lepsch, L.B., Gonzalo, L.A., Magro, F.J.B., Delucia, R., Scavone, C., Planeta, C.S., 2005. Exposure to chronic stress increases the locomotor response to cocaine and the basal levels of corticosterone in adolescent rats. Addict. Biol, 10(3), 251–256. 10.1080/13556210500269366

Li, Y., Hu, X., Berney, T.G., Vartanian, A.J., Stine, C.D., Wolf, M.E., White, F.J., 1999. Both glutamate receptor antagonists and prefrontal cortex lesions prevent induction of cocaine sensitization and associated neuroadaptations. Synapse, 34(3), 169–180. 10.1002/(sici)1098-2396(19991201)34:3<169::aid-syn1>3.3.co;2-3

Lohani, S., Martig, A.K., Deisseroth, K., Witten, I.B., Moghaddam, B., 2019. Dopamine Modulation of Prefrontal Cortex Activity Is Manifold and Operates at Multiple Temporal and Spatial Scales. Cell Rep, 27(1), 99–114.e6. 10.1016/j.celrep.2019.03.012

Long, S.F., Dennis, L.A., Russell, R.K., Benson, K.A., Wilson, M.C., 1994. Testosterone implantation reduces the motor effects of cocaine. Behav. Pharmacol, 5(1), 103. 10.1097/00008877-199402000-00012

Lood, Y., Eklund, A., Garle, M., Ahlner, J., 2012. Anabolic androgenic steroids in police cases in Sweden 1999-2009. Forensic Sci. Int. 219, 199–204. 10.1016/j.forsciint.2012.01.004

Lukas, S.E., Sholar, M., Lundahl, L.H., Lamas, X., Kouri, K., Wines, J.D., Kragie, L., Mendelson, J.H., 1996. Sex differences in plasma cocaine levels and subjective effects after acute cocaine administration in human volunteers. Psychopharmacology (Berl), 125(4), 346–354. 10.1007/BF02246017

Malone, D.A., Dimeff, R.J., Lombardo, J.A., Sample, R.H.B., 1995. Psychiatric effects and psychoactive substance use in anabolic-androgenic steroid users. Clin. J. Sport Med, 5(1), 25–31. 10.1097/00042752-199501000-00005

Manvich, D.F., Petko, A.K., Branco, R.C., Foster, S.L., Porter-Stransky, K.A., Stout, K.A., Newman, A.H., Miller, G.W., Paladini, C.A., Weinshenker, D., 2019. Selective D2 and D3 receptor antagonists oppositely modulate cocaine responses in mice via distinct postsynaptic mechanisms in nucleus accumbens. Neuropsychopharmacology, 44(8), 1445–1455. 10.1038/s41386-019-0371-2

Marinelli, M., White, F.J., 2000. Enhanced vulnerability to cocaine self-administration is associated with elevated impulse activity of midbrain dopamine neurons. J. Neurosci, 20(23), 8876–8885. 10.1523/jneurosci.20-23-08876.2000

Mattingly, B.A., Hart, T.C., Lim, K., Perkins, C., 1994. Selective antagonism of dopamine D1 and D2 receptors does not block the development of behavioral sensitization to cocaine. Psychopharmacology (Berl). 10.1007/BF02244843

McDonnell, A. M., & Dang, C. H., 2013. Basic review of the cytochrome p450 system. Journal of the advanced practitioner in oncology, 114(2), 239–242. ttps://doi.org/10.6004/jadpro.2013.4.4.7

McGinty, J.F., Whitfield, T.W., Berglind, W.J., 2010. Brain-derived neurotrophic factor and cocaine addiction. Brain Res, 1314, 183–193. 10.1016/j.brainres.2009.08.078

Mȩdraś, M., Brona, A., Jóźków, P., 2018. The Central Effects of Androgenic-Anabolic Steroid Use. J. Addict. Med, 12(3), 184–192. 10.1097/ADM.0000000000000395

Menéndez-Delmestre, R., Segarra, A.C., 2011. Testosterone is essential for cocaine sensitization in male rats. Physiol. Behav, 102(1), 96–104. 10.1016/j.physbeh.2010.09.025

Metastasio, A., Negri, A., Martinotti, G., Corazza, O., 2018. Transitioning bodies. The case of self-prescribing sexual hormones in gender affirmation in individuals attending psychiatric services. Brain Sci, 8(5), 88. 10.3390/brainsci8050088

Minkin, D.M., Meyer, M.E., van Haaren, F., 1993. Behavioral effects of long-term administration of an anabolic steroid in intact and castrated male Wistar rats. Pharmacol. Biochem. Behav, 44(4), 959–963. 10.1016/0091-3057(93)90031-N

Moorman, D.E., James, M.H., McGlinchey, E.M., Aston-Jones, G., 2015. Differential roles of medial prefrontal subregions in the regulation of drug seeking. Brain Res, 1628, 130–146. 10.1016/j.brainres.2014.12.024

Morrison, T.R., Ricci, L.A., Melloni, R.H., 2015. Aggression and anxiety in adolescent AAS-treated hamsters: A role for 5HT _3_ receptors. Pharmacol. Biochem. Behav. 134, 85–91. 10.1016/j.pbb.2015.05.001

Mueller, D., Chapman, C.A., Stewart, J., 2006. Amphetamine induces dendritic growth in ventral tegmental area dopaminergic neurons in vivo via basic fibroblast growth factor. Neuroscience, 1628, 130–146. 10.1016/j.neuroscience.2005.09.038

Neisewander, J.L., Fuchs, R.A., O’Dell, L.E., Khroyan, T. V., 1998. Effects of SCH-23390 on dopamine D1 receptor occupancy and locomotion produced by intraaccumbens cocaine infusion. Synapse, 30(2), 194–204. 10.1002/(sici)1098-2396(199810)30:2<194::aid-syn9>3.3.co;2-a

Novoa, J., Rivero, C.J., Pérez-Cardona, E.U., Freire-Arvelo, J.A., Zegers, J., Yarur, H.E., Santiago-Marerro, I.G., Agosto-Rivera, J.L., González-Pérez, J.L., Gysling, K., Segarra, A.C., 2021. Social isolation of adolescent male rats increases anxiety and K+-induced dopamine release in the nucleus accumbens: Role of CRF-R1. Eur. J. Neurosci, 54(3), 4888–4905. 10.1111/ejn.15345

Oberlander, J.G., Henderson, L.P., 2012. The Sturm und Drang of anabolic steroid use: Angst, anxiety, and aggression. Trends Neurosci, 35(6), 382–392. 10.1016/j.tins.2012.03.001

Onakomaiya, M.M., Henderson, L.P., 2016. Mad men, women and steroid cocktails: A review of the impact of sex and other factors on anabolic androgenic steroids effects on affective behaviors. Psychopharmacology (Berl), 233(4), 549–569. 10.1007/s00213-015-4193-6

Op De MacKs, Z.A., Moor, B.G., Overgaauw, S., Gürolu, B., Dahl, R.E., Crone, E.A., 2011. Testosterone levels correspond with increased ventral striatum activation in response to monetary rewards in adolescents. Dev. Cogn. Neurosci, 1(4), 506–516. 10.1016/j.dcn.2011.06.003

Pagonis, T.A., Angelopoulos, N. V., Koukoulis, G.N., Hadjichristodoulou, C.S., 2006. Psychiatric side effects induced by supraphysiological doses of combinations of anabolic steroids correlate to the severity of abuse. Eur. Psychiatry, 21(8), 551–562. 10.1016/j.eurpsy.2005.09.001

Parkinson, A.B., Evans, N.A., 2006. Anabolic androgenic steroids: A survey of 500 users. Med. Sci. Sports Exerc, 38(4), 644–651. 10.1249/01.mss.0000210194.56834.5d

Paus, T., Keshavan, M., Giedd, J.N., 2008. Why do many psychiatric disorders emerge during adolescence? Nat. Rev. Neurosci, 9(12), 947–957. 10.1038/nrn2513

Pellow, S., Chopin, P., File, S.E., Briley, M., 1985. Validation of open : closed arm entries in an elevated plus-maze as a measure of anxiety in the rat. J. Neurosci. Methods, 14(3), 149–167. 10.1016/0165-0270(85)90031-7

Penatti, C.A.A., Oberlander, J.G., Davis, M.C., Porter, D.M., Henderson, L.P., 2011. Chronic exposure to anabolic androgenic steroids alters activity and synaptic function in neuroendocrine control regions of the female mouse. Neuropharmacology, 61(4), 653–664. 10.1016/j.neuropharm.2011.05.008

Peters, K.D., Wood, R.I., 2005. Androgen dependence in hamsters: Overdose, tolerance, and potential opioidergic mechanisms. Neuroscience, 130(4), 971–981. 10.1016/j.neuroscience.2004.09.063

Pope, H.G., Kanayama, G., Athey, A., Ryan, E., Hudson, J.I., Baggish, A., 2014a. The lifetime prevalence of anabolic-androgenic steroid use and dependence in Americans: Current best estimates. Am. J. Addict, 23(4), 371–377. 10.1111/j.1521-0391.2013.12118.x

Pope, H.G., Katz, D.L., 1988. Affective and psychotic symptoms associated with anabolic steroid use. Am. J. Psychiatry, 145(4), 487–490. 10.1176/ajp.145.4.487

Pope, H.G., Kouri, E.M., Powell, K.F., Campbell, C., Katz, D.L., 1996. Anabolic-androgenic steroid use among 133 prisoners. Compr. Psychiatry, 37(5), 322–327. 10.1016/S0010-440X(96)90013-9

Pope, H.G., Wood, R.I., Rogol, A., Nyberg, F., Bowers, L., Bhasin, S., 2014b. Adverse health consequences of performance-enhancing drugs: An endocrine society scientific statement. Endocr. Rev, 35(3), 341–375. 10.1210/er.2013-1058

Purves-Tyson, T.D., Allen, K., Fung, S., Rothmond, D., Noble, P.L., Handelsman, D.J., Shannon Weickert, C., 2015. Adolescent testosterone influences BDNF and TrkB mRNA and neurotrophin-interneuron marker relationships in mammalian frontal cortex. Schizophr. Res, 168(3), 661–670. 10.1016/j.schres.2015.05.040

Purves-Tyson, T.D., Handelsman, D.J., Double, K.L., Owens, S.J., Bustamante, S., Weickert, C.S., 2012. Testosterone regulation of sex steroid-related mRNAs and dopamine-related mRNAs in adolescent male rat substantia nigra. BMC Neurosci, 13(1), 95. 10.1186/1471-2202-13-95

Rainer Q, Spezial, S, Rubino T, Dominguez-Lopez S, Rodriguez Bambico F, Gobbi G and D Parolaro. 2014. Chronic nandrolone decanoate exposure during adolescence affects emotional behavior and monoaminergic neurotransmission in adulthood. Neuropharmacology, v. 83, p. 79-88, 2014. doi: 10.1016/j.neuropharm.2014.03.015. Epub 2014 Apr 8.

Ricci, L.A., Morrison, T.R., Melloni, R.H., 2012. Serotonin modulates anxiety-like behaviors during withdrawal from adolescent anabolic-androgenic steroid exposure in Syrian hamsters. Horm. Behav. 62, 569–578. 10.1016/j.yhbeh.2012.09.007

Rice, M.E., Cragg, S.J., 2008. Dopamine spillover after quantal release: Rethinking dopamine transmission in the nigrostriatal pathway. Brain Res. Rev, 58(2), 303–313. 10.1016/j.brainresrev.2008.02.004

Riedel, W.J., Klaassen, T., Deutz, N.E.P., Van Someren, A., Van Praag, H.M., 1999. Tryptophan depletion in normal volunteers produces selective impairment in memory consolidation. Psychopharmacology (Berl), 141(4), 362–369. 10.1007/s002130050845

Robinson, T.E., Berridge, K.C., 1993. The neural basis of drug craving: An incentive-sensitization theory of addiction. Brain Res. Rev, 18(3), 247–291. 10.1016/0165-0173(93)90013-P

Rosic, G., Joksimovic, J., Selakovic, D., Milovanovic, D., Jakovljevic, V., 2014. Anxiogenic effects of chronic exposure to nandrolone decanoate (ND) at supraphysiological dose in rats: A brief report. Neuroendocrinol. Lett, 35(8), 703– 710.

Saartok, T., Dahlberg, E., Gustafsson, J.Å., 1984. Relative binding affinity of anabolic-androgenic steroids: Comparison of the binding to the androgen receptors in skeletal muscle and in prostate, as well as to sex hormone-binding globulin. Endocrinology, 114(6), 2100–2106. 10.1210/endo-114-6-2100

Sagoe, D., Molde, H., Andreassen, C.S., Torsheim, T., Pallesen, S., 2014. The global epidemiology of anabolic-androgenic steroid use: A meta-analysis and meta-regression analysis. Ann. Epidemiol, 24(5), 383–398. 10.1016/j.annepidem.2014.01.009

Saito, H., Kasayama, S., Kouhara, H., Matsumoto, K., Sato, B., 1991. Up-regulation of fibroblast growth factor (FGF) receptor mRNA levels by basic FGF or testosterone in androgen-sensitive mouse mammary tumor cells. Biochem. Biophys. Res. Commun, 174(1), 136–141. 10.1016/0006-291X(91)90496-T

Salas, J., Scherrer, J.F., Lustman, P.J., Schneider, F.D., 2016. Racial differences in the association between nonmedical prescription opioid use, abuse/dependence, and major depression. Subst. Abus. 37, 25–30. 10.1080/08897077.2015.1129523

Salvador, A., Moya-Albiol, L., Martínez-Sanchis, S., Simón, V.M., 1999. Lack of effects of anabolic-androgenic steroids on locomotor activity in intact male mice. Percept. Mot. Skills, 88(1), 319–328. 10.2466/pms.1999.88.1.319

SAMHSA, 2019. Substance Abuse and Mental Health Services Administration, results from the 2018 National Survey on Drug Use and Health: Detailed tables. Rockville, MD: Center for Behavioral Health Statistics and Quality, Substance Abuse and Mental Health Services Administration.

Segarra, A.C., Strand, F.L., 1989. Perinatal administration of nicotine alters subsequent sexual behavior and testosterone levels of male rats. Brain Res, 480(1–2), 151–159. 10.1016/0006-8993(89)91577-1

Segarra AC, Torres-Díaz YM, Silva RD, Puig-Ramos A, Menéndez-Delmestre R, Rivera-Bermúdez JG, Amadeo W, Agosto-Rivera JL 2014. Estrogen receptors mediate estradiol’s effect on sensitization and CPP to cocaine in female rats: Role of contextual cues. Horm Behav 65 (2): 77–87. doi: 10.1016/j.yhbeh.2013.12.007. Epub 2013 Dec 17.

Selakovic, D., Joksimovic, J., Zaletel, I., Puskas, N., Matovic, M., Rosic, G., 2017. The opposite effects of nandrolone decanoate and exercise on anxiety levels in rats may involve alterations in hippocampal parvalbumin-positive interneurons. PLoS One, 12(12), e0189595. 10.1371/journal.pone.0189595

Shinohara, F., Kamii, H., Minami, M., Kaneda, K., 2017. The role of dopaminergic signaling in the medial prefrontal cortex for the expression of cocaine-induced conditioned place preference in rats. Biol. Pharm. Bull, 40(11), 1983–1989. 10.1248/bpb.b17-00614

Sjöqvist, F., Garle, M., Rane, A., 2008. Use of doping agents, particularly anabolic steroids, in sports and society. Lancet, 371(9627), 1872–1882. 10.1016/S0140-6736(08)60801-6

Smith, R.J., Lobo, M.K., Spencer, S., Kalivas, P.W., 2013. Cocaine-induced adaptations in D1 and D2 accumbens projection neurons (a dichotomy not necessarily synonymous with direct and indirect pathways). Curr. Opin. Neurobiol, 23(4), 546–552. 10.1016/j.conb.2013.01.026

Steketee, J.D., Kalivas, P.W., 2011. Drug wanting: Behavioral sensitization and relapse to drug-seeking behavior. Pharmacol. Rev, 63(2), 348–365. 10.1124/pr.109.001933

Stopper, C.M., Floresco, S.B., 2015. Dopaminergic circuitry and risk/reward decision making: Implications for schizophrenia. Schizophr. Bull, 41(1), 9–14. 10.1093/schbul/sbu165

Stopper, C.M., Khayambashi, S., Floresco, S.B., 2013. Receptor-specific modulation of risk-based decision making by nucleus accumbens dopamine. Neuropsychopharmacology, 38(5), 715–728. 10.1038/npp.2012.240

Tenniswood, M., Bird, C.E., Clark, A.F., 1982. The role of androgen metabolism in the control of androgen action in the rat prostate. Mol. Cell. Endocrinol, 27(1), 89–96. 10.1016/0303-7207(82)90065-X

Thomas, M.J., Beurrier, C., Bonci, A., Malenka, R.C., 2001. Long-term depression in the nucleus accumbens: A neural correlate of behavioral sensitization to cocaine. Nat. Neurosci, 4(12), 1217–1223. 10.1038/nn757

Thomas, M.J., Kalivas, P.W., Shaham, Y., 2008. Neuroplasticity in the mesolimbic dopamine system and cocaine addiction. Br. J. Pharmacol, 154(2), 327–342. 10.1038/bjp.2008.77

Tobiansky, D.J., Wallin-Miller, K.G., Floresco, S.B., Wood, R.I., Soma, K.K., 2018. Androgen regulation of the mesocorticolimbic system and executive function. Front. Endocrinol, 9.10.3389/fendo.2018.00279

Treit, D., Fundytus, M., 1988. Thigmotaxis as a test for anxiolytic activity in rats. Pharmacol. Biochem. Behav. 31(4), 959–962. 10.1016/0091-3057(88)90413-3

Trzcińska, M., Bergh, J., DeLeon, K., Stellar, J.R., Melloni, R.H., 2002. Social stress does not alter the expression of sensitization to cocaine. Physiol. Behav, 76(4–5), 457–463. 10.1016/S0031-9384(02)00727-8

Tseng, Y,T,, Rockhold, R.W., Hoskins, B., Ho, I.K. 1994. Cardiovascular toxicities of nandrolone and cocaine in spontaneously hypertensive rats. Fundam Appl Toxicil 22(1): 113–121. 10.1006/faat.1994.1014.

Turner, C.A., Capriles, N., Flagel, S.B., Perez, J.A., Clinton, S.M., Watson, S.J., Akil, H., 2009. Neonatal FGF2 alters cocaine self-administration in the adult rat. Pharmacol. Biochem. Behav, 92(1), 100–104. 10.1016/j.pbb.2008.10.018

van Amsterdam, J., Opperhuizen, A., Hartgens, F., 2010. Adverse health effects of anabolic-androgenic steroids. Regul. Toxicol. Pharmacol, 57(1), 117–123. 10.1016/j.yrtph.2010.02.001

Vanderschuren, L.J.M.J., Kalivas, P.W., 2000. Alterations in dopaminergic and glutamatergic transmission in the induction and expression of behavioral sensitization: A critical review of preclinical studies. Psychopharmacology (Berl), 151(2–3), 99–120. 10.1007/s002130000493

Vanderschuren, L.J.M.J., Pierce, R.C., 2010. Sensitization processes in drug addiction. Curr. Top. Behav. Neurosci, (pp. 179–195). 10.1007/7854_2009_21

Van der Vies, J., 1985. Implications of basic pharmacology in the therapy with esters of nandrolone. Acta Endocrinol. Suppl. 110(3_Suppla), S38–S44. 10.1530/acta.0.109s0038

Wallin-Miller, K.G., Kreutz, F., Li, G., Wood, R.I., 2018. Anabolic-androgenic steroids (AAS) increase sensitivity to uncertainty by inhibition of dopamine D1 and D2 receptors. Psychopharmacology (Berl). 235, 959–969. 10.1007/s00213-017-4810-7.

Wallin, K.G., Alves, J.M., Wood, R.I., 2015. Anabolic-androgenic steroids and decision making: Probability and effort discounting in male rats. Psychoneuroendocrinology, 57, 84–92. 10.1016/j.psyneuen.2015.03.023

Wallin, K.G., Wood, R.I., 2015. Anabolic-androgenic steroids impair set-shifting and reversal learning in male rats. Eur. Neuropsychopharmacol, 25(4), 583–590. 10.1016/j.euroneuro.2015.01.002

White, F.J., Joshi, A., Koeltzow, T.E., Hu, X.T., 1998. Dopamine receptor antagonists fail to prevent induction of cocaine sensitization. Neuropsychopharmacology, 18(1), 26–40. 10.1016/S0893-133X(97)00093-6

Winstanley, C.A., Floresco, S.B., 2016. Deciphering decision making: Variation in animal models of effort- and uncertainty-based choice reveals distinct neural circuitries underlying core cognitive processes. J. Neurosci, 36(48), 12069–12079. 10.1523/JNEUROSCI.1713-16.2016

Wood, R.I., 2004. Reinforcing aspects of androgens. Physiol. Behav, 83(2), 279–289. 10.1016/j.physbeh.2004.08.012

Wood, R.I., 2002. Oral testosterone self-administration in male hamsters: Dose-response, voluntary exercise, and individual differences. Horm. Behav, 41(3), 247–258. 10.1006/hbeh.2002.1769

Wood, R.I., Armstrong, A., Fridkin, V., Shah, V., Najafi, A., Jakowec, M., 2013. ’Roid rage in rats? Testosterone effects on aggressive motivation, impulsivity and tyrosine hydroxylase. Physiol. Behav, 110–111, 6–12. 10.1016/j.physbeh.2012.12.005

Yamamoto, R.T., Teter, C.J., Barros, T.L., McCarthy, E., Mileti, C., Juliano, T., Medeiros, C.L., Looby, A., Maywalt, M.A., McNeil, J.F., Olson, D., Mallya, G., Lukas, S.E., Renshaw, P.F., Kaufman, M.J., 2007. Antiandrogen pretreatment alters cocaine pharmacokinetics in men. J. Addict. Med, 1(4), 198–204. 10.1097/ADM.0b013e31815a137c

Yokel, R.A., Wise, R.A., 1975. Increased lever pressing for amphetamine after pimozide in rats: Implications for a dopamine theory of reward. Science, 187(4176), 547–549. 10.1126/science.1114313

Zotti, M., Tucci, P., Colaianna, M., Morgese, M.G., Mhillaj, E., Schiavone, S., Scaccianoce, S., Cuomo, V., Trabace, L., 2014. Chronic nandrolone administration induces dysfunction of the reward pathway in rats. Steroids 79, 7–13. 10.1016/j.steroids.2013.10.005

